# Fuzzy Supertertiary Interactions within PSD-95 Enable Ligand Binding

**DOI:** 10.1101/2022.02.23.481591

**Authors:** George L. Hamilton, Nabanita Saikia, Sujit Basak, Franceine S. Welcome, Fang Wu, Jakub Kubiak, Changcheng Zhang, Yan Hao, Claus A.M. Seidel, Feng Ding, Hugo Sanabria, Mark E. Bowen

## Abstract

The scaffold protein PSD-95 links postsynaptic receptors to sites of presynaptic neurotransmitter release. Flexible linkers between folded domains in PSD-95 enable a dynamic supertertiary structure. Interdomain interactions within the PSG supramodule, formed by PDZ3, SH3 and GuK domains, regulate PSD-95 activity. Here we combined Discrete Molecular Dynamics and single molecule FRET to characterize the PSG supramodule, with time resolution spanning picoseconds to seconds. We used a FRET network to measure distances in full-length PSD-95 and model the conformational ensemble. We found that PDZ3 samples two conformational basins, which we confirmed with disulfide mapping. To understand effects on activity, we measured binding of the synaptic adhesion protein neuroligin. We found that PSD-95 bound neuroligin well at physiological pH while truncated PDZ3 bound poorly. Our hybrid structural models reveal how the supertertiary context of PDZ3 enables recognition of this critical synaptic ligand.

## INTRODUCTION

The fundamental structural unit of large proteins is the independently folding domain ^1^. Most proteins are composed of more than one domain ^2,3^. Evolution has shuffled the deck to produce a rich diversity of multidomain proteins. A prototypical example is the Membrane Associated Guanylate Kinases (MAGuKs), which contain an array of folded protein-interaction domains connected in series. MAGuKs are scaffold proteins that link cell surface membrane proteins to their intracellular signaling partners and the cytoskeleton ^4^. MAGuK family members control diverse processes ranging from epithelial cell polarity ^5^ to synaptic neurotransmitter signaling ^6^.

Almost all MAGuKs contain a conserved “PSG” supramodule that links a PDZ domain ^7^ an SH3 domain ^8^ and guanylate kinase-like (GuK) domain ^9,10^ that serves as another protein-binding domain ^11^ (Figure 1A). One distinctive feature of the MAGuKs is the insertion of a variable HOOK region within the SH3 domain that disrupts canonical SH3 domain interactions ^12^. Different MAGuKs append additional protein-binding domains to this PSG supramodule ^5^. The postsynaptic density protein of 95 kilodaltons (PSD-95) contains an N-terminal extension and two tandem PDZ domains attached to its PSG (Figure 1A).

**Figure 1.**
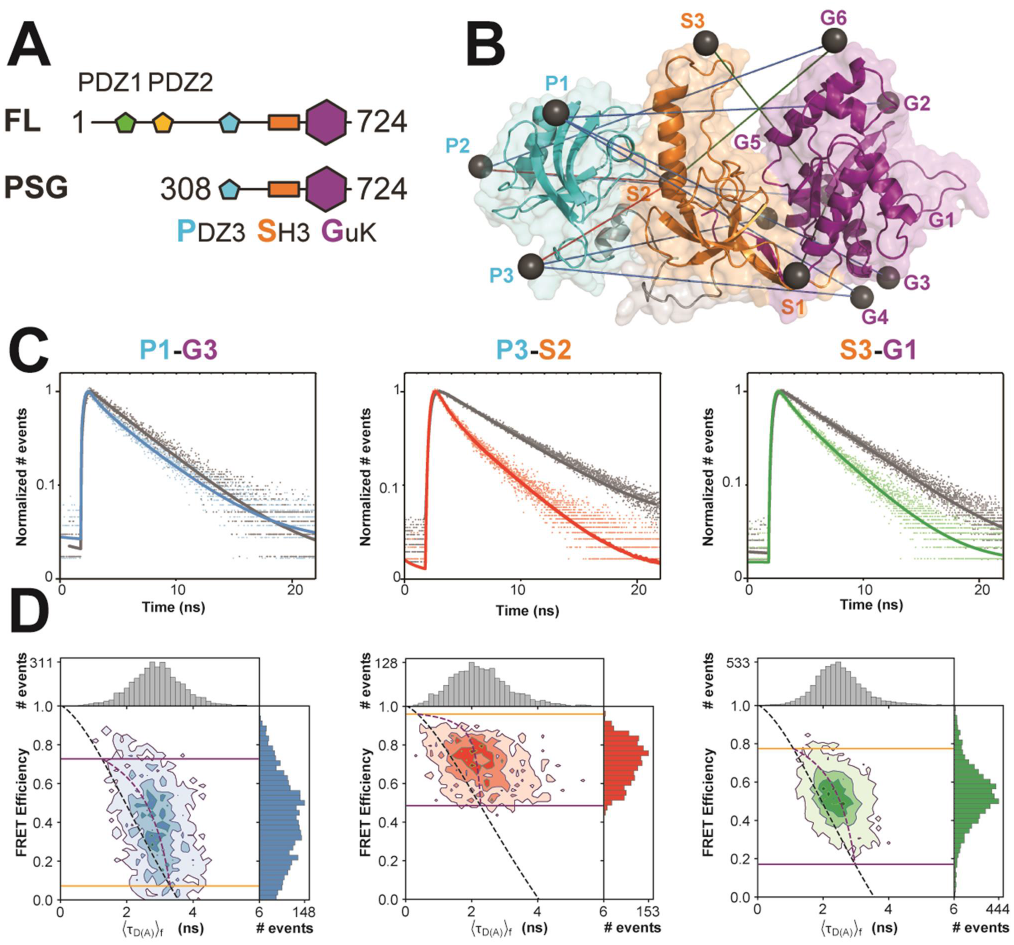
Multiparameter Fluorescence Detection of Single-Molecule FRET in PSD-95. **A)** Cartoon representation of domain organization in PSD-95. **B)** Location of the cysteine substitutions used for fluorescent labeling. Spheres indicate the location of the dye while lines indicate the experimental FRET pairs. Details about labeling sites can be found in Table 1. Although only the PSG is shown, data shown is from measurements of full-length PSD-95. **C)** Representative fluorescence decays from seTCSPC. Shown are Donor-Only controls (grey) and matched FRET samples for variants P1-G3 (blue), P3-S2 (red) and S3-G1 (green). The FRET decays were fit using a global analysis of all 12 variants. **D)** Representative 2D MFD histograms for the variants shown in panel C. Fluorescence lifetime (x-axis) is plotted against the FRET efficiency for each molecule. The seTCSPC limiting states A (orange) and B (purple) are shown as solid lines. The expected relationship between fluorescence lifetime and FRET is shown for static (black dashes) and dynamic molecules (purple dashes).

In multidomain proteins, the proximity imparted by interdomain linkers results in high effective concentrations of the folded domains ^13^. This can result in specific domain interactions that are too weak to manifest when the domains are not connected ^14^. Linkers also allow for different supertertiary arrangements that must be represented by a multiplicity of states ^15^. The presence of heterogeneity confounds structural biology methods reliant on ensemble averaging while protein dynamics leads to time averaging even in most single-molecule experiments ^16^.

Mapping the conformational landscape of the PSG is important because these supertertiary interactions regulate protein interactions with scaffolding clients ^17-19^. Previous SAXS and NMR studies along with computational approaches have proposed binding sites for PDZ3 within SH3-GuK (or a lack thereof) ^20-22^. Single-molecule Förster resonance energy transfer (smFRET) studies revealed that PDZ3 was dynamic but had a defined orientation relative to SH3-GuK ^22^.

Recently, we combined multiparameter fluorescence measurements with discrete molecular dynamics (DMD) and disulfide mapping to characterize the PDZ tandem from PSD-95 ^23^. Here, we applied this approach to the PSG supramodule. Experiments and simulations agreed and revealed two distinct conformational basins for PDZ3; a fuzzy interaction with PDZ3 within a broad interface near the SH3 HOOK insertion and a second discrete binding site in GuK. Both were confirmed with disulfide mapping. Surprisingly, these supertertiary interactions in full-length PSD-95 allowed PDZ3 to interact with the synaptic adhesion protein neuroligin, a known binding partner of PDZ3 ^24-26^, while the truncated domain showed weaker binding. Thus, the supertertiary context enhanced the binding activity of PDZ3 towards a critical physiological ligand.

Our integrative approach for dynamic structural biology resolved the structural heterogeneity of the PSG supramodule within full-length PSD-95. Combining simulation with experiments resolved global dynamics and provided residue-level details about supertertiary interactions. This approach moves beyond solving a single structure towards resolving an ensemble of conformers with differing behavior and is applicable to other dynamic multidomain proteins.

## RESULTS

### Mapping the Supertertiary Conformational Landscape with Single-Molecule FRET

To experimentally probe the location of PDZ3 within the PSG supramodule (Figure 1A), we used 11 cysteine mutations spanning PDZ3, SH3, and GuK (Table 1). By using these labeling sites in different combinations, we created 12 variants (Figure 1B), which form a FRET network for structural modeling (Figure 1 – figure supplement 1) ^16^. Fluorescently labeled proteins were measured with confocal multiparameter fluorescence detection (MFD; for instrument and correction parameters see Supplementary file 1A &B) and pulsed-interleaved excitation (PIE) that allowed selection of molecules with active donor and acceptor fluorophores for sub-ensemble Time-Correlated Single Photon Counting (seTCSPC) (Figure 1C & Figure 1 – figure supplement 2) ^27^. Each variant provides specific information on the same conformational landscape. To capture this shared information, we performed a simultaneous global analysis of all variants ^28^. The FRET distances were variant-specific while the number and occupancy of conformational states were set as global fitting parameters. This assumes that the population distribution is the same for all variants, but that each variant senses the underlying conformations differently. Based on fitting statistics, we demonstrate that a two-state model with a small donor-only (or no FRET) population (Supplementary file 1C &D) is sufficient to fit all data. Thus, PDZ3 samples two limiting states with slight predominance of state B (53.9%) over state A (46.1%). The global fit assigns all distances to their respective states (Supplementary file 1E).

**Table 1.** Labeling sites used for single-molecule FRET measurements. The labeling position is identified first by a single letter indicating the domain (P for PDZ3, S for SH3 and G for GuK) followed by its sequential order in the primary sequence.

| Position | Mutation | **Table 1. Labeling sites used for single-molecule FRET measurements.** The labeling position is identified first by a single letter indicating the domain (P for PDZ3, S for SH3 and G for GuK) followed by its sequential order in the primary sequence. |
| --- | --- | --- |
| P1 | R313C |  |
| P2 | Q374C |  |
| P3 | S398C |  |
| S1 | H477C |  |
| S2 | R492C |  |
| S3 | S510C |  |
| G1 | K591C |  |
| G2 | S606C |  |
| G3 | E618C |  |
| G4 | E621C |  |
| G5 | A640C |  |
| G6 | R671C |  |

For each variant, we plotted the intensity-based FRET efficiency against the average donor fluorescence lifetime (⟨*τ*_*D*(*A*)_⟩_*f*_) for each molecule (Figure 1D & Figure 1 – figure supplement 3). We calculated the expected relationship between FRET and ⟨*τ*_*D*(*A*)_⟩_*f*_ using the assumption of no dynamics, which we plotted as the static FRET-line ^29^ (Supplementary file 1F). Molecules undergoing dynamics would fall off this line. The dynamic FRET-lines represent all possible degrees of mixing between the limiting states (Supplementary file 1G). All measured variants exhibited a rightward skew away from the static FRET-line, a hallmark of conformational dynamics (Figure 1D & Figure 1 – figure supplement 3) ^29^.

Variants involving PDZ3 exhibited broad or irregular distributions indicating a heterogeneous conformational ensemble. Variants between PDZ3 and SH3 exhibited higher FRET efficiency, suggesting close proximity. The SH3-GuK domains have restricted interdomain motion ^12,30^. Nonetheless, FRET variants spanning SH3-GuK still fell off the static FRET-line. These variants exhibited narrower, more regular FRET distributions relative to PDZ3-labeled variants indicating fast but limited dynamics within SH3-GuK.

### Comparison of full-length PSD-95 to the PSG truncation

To probe whether interactions within PSD-95 affect the PSG, we also measured 6 of the FRET variants in a truncated PSG fragment. Measurements using smTIRF with camera detection revealed changes in the time-averaged FRET distributions for all variants (Figure 2A & Figure 2 – figure supplement 1). Truncating PSD-95 resulted in broader and more multi-peaked distributions. For example, variant P1-G3 (Table 1), splits into lower and higher FRET in the truncation. (Figure 2A). Similarly, variant P3-S2 showed the highest FRET but the distribution spread out to lower FRET when PSD-95 is truncated. Truncation also increased anticorrelated FRET transitions in smTIRF time traces suggesting altered dynamics of PDZ3 (Figure 2 – figure supplement 1C). To quantify this, we determined the donor-acceptor cross-correlation amplitude. While the magnitude of the cross-correlation amplitudes depends on FRET efficiency and rate constants, comparison of the same labeling sites in full-length and PSG revealed a uniform increase in FRET transitions when PSD-95 is truncated (Figure 2 – figure supplement 1D).

**Figure 2.**
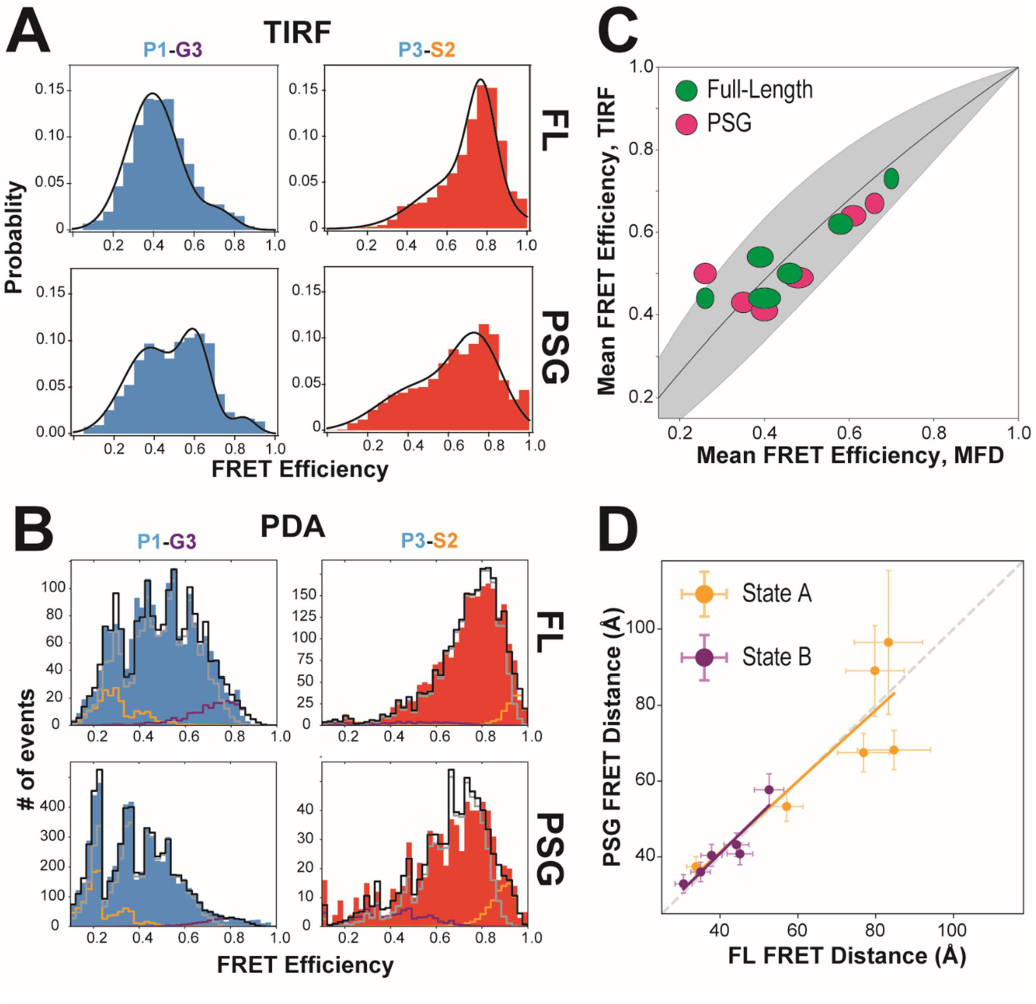
Effect of Truncating PSD-95 on PSG Supertertiary Structure. **A)** Representative smTIRF FRET efficiency histograms for full-length PSD-95 (top) and the corresponding truncated PSG fragment (bottom). Shown are variants P1-G3 (blue) and P3-S2 (red). **B)** Representative PDA plots using a 2 ms time window for the same variants from panel A using the same coloring. Molecules occupying limiting states A and B are highlighted in orange and purple, respectively. **C)** Comparison of mean FRET as measured with smTIRF (y-axis) and MFD (x-axis) for full-length (green) and PSG (pink) variants. Ellipse eccentricities represent the relative width of FRET distributions observed by each method. The expected relationship given the different fluorophores used is shown as a line with the shaded region corresponding to Förster Radius uncertainty. Förster Radii used were as used previously ^22,23^. The fit to the ideal relationship gave 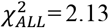 with 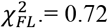 and 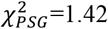. **D)** Comparison of the seTCSPC limiting-state distances for full-length PSD-95 (y-axis) and the PSG fragment (x-axis). Distances are shown for state A (orange; Slope =0.94; Pearson Correlation Coefficient (*R*_*p*_) = 0.86) and state B (purple; Slope =1.0; *R*_*p*_ = 0.93).

Measurements with MFD revealed shifts in FRET efficiency and ⟨*τ*_*D*(*A*)_⟩_*f*_ for most variants (Figure 2 – figure supplement 2). To further analyze the intensity-based FRET efficiencies, we performed Photon Distribution Analysis (PDA) ^31^. Our PDA fit model included the two static, limiting states and a dynamic population (Supplementary file 2). Several truncated variants had more molecules in static limiting states, indicating increased dwell time of PDZ3 (Figure 2 – figure supplement 3). This suggests the slowest exchange processes were further slowed in truncated variants. However, the predominant state for PDZ3 was always in exchange with faster relaxation rates. PDA additionally found truncation-induced shifts in FRET efficiency similar to smTIRF (Figure 2B). The good agreement between mean FRET efficiencies measured with smTIRF and MFD, representing the long-time averages from both techniques, brings additional confidence in the results (Figure 2C). A global fit of seTCSPC for the PSG recovered two states similar to the full-length protein (Figure 2D & Figure 2 – figure supplement 4). Truncating PSD-95 shifted the limiting state distances for state A and slightly reduced state B occupancy (48.2%, Supplementary file 1E).

To resolve fast conformational dynamics, we performed filtered fluorescence correlation spectroscopy (fFCS) ^32^. We filtered bursts into subensembles representing the seTCSPC limiting states and analyzed these components using standard correlation algorithms ^32^ (Figure 3 – figure supplement 1). Just as FRET efficiency differed between variants, each variant is differently attuned to the same underlying conformational transitions, so data was fit globally to capture the shared information (results in Supplementary file 3A & B). Three decay times were assigned to local motions (*t*_*R1*_) that maintain residue contacts, domain re-orientations (*t*_*R2*_) that alter interdomain interaction interfaces, and domain exchange (*t*_*R3*_) such as large-scale translational transitions between basins (Figure 3A). PSD-95 variants displayed complex dynamics with components from µs to ms. To highlight differences between variants, we plotted the normalized relaxation amplitudes in a matrix representation, which is the dynamics equivalent of a contact map for protein interactions (Figure 3B). This revealed that 7 out of the 10 GuK-labeled variants have major (red) or middle (yellow) contributions at *t*_*R1*_. The remaining 3 variants are dominated by *t*_*R3*_. The large contribution at *t*_*R1*_ for most PDZ3-GuK variants suggests fast local motions within the limiting states, while for P2-G5 and P3-G5 domain exchange dictated the dynamics. We also note that for 4 of the remaining 5 PDZ3-GuK variants, the *t*_*R3*_ has middle or major contributions. Moreover, P2-G5, P3-G4, and P3-G5 reported middle contributions from domain reorientation.

**Figure 3.**
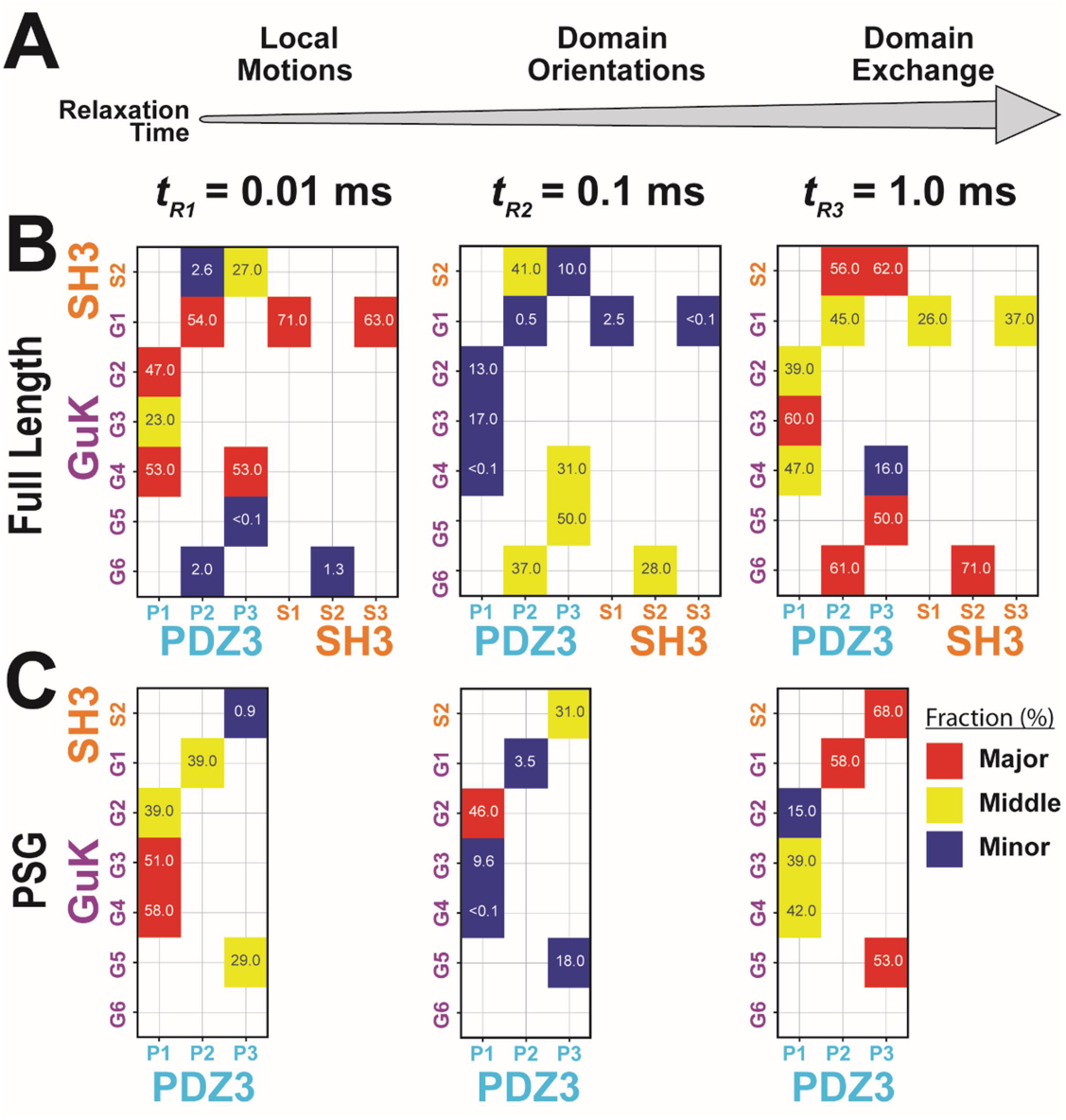
Effect of Truncating PSD-95 on Supertertiary Dynamics within PSG. Site-specific dynamics for each variant as measured with Filtered Fluorescence Correlation Spectroscopy (fFCS) and fit to a model with three correlation times at three decades in time from 10^−5^ to 10^−3^ seconds. **A)** Association between the dynamic relaxation time and the expected protein motions across the three decades in time used in the analysis. **B)** Matrix representation of the relative contribution to dynamic relaxation in full length PSD-95 at each decade in time (as indicated above the panel). The axes specify the domains and labeling sites with each variant placed at the intersection between sites used. The major timescale for relaxation is highlighted in red; the minor in blue and the middle in yellow with the percentage within each square. **C)** Matrix representation of dynamics in the PSG truncation measured with the same variants and shown with coloring identical to panel B. The major populations for both full length and PSG variants are mapped to the sequence and secondary structure in Figure 3 - figure supplement 2.

Summarizing the dynamics observed for the PDZ3-GuK variants, fFCS depicts three relaxation times. The major contributions are either *t*_*R1*_ suggesting fast domain motions within identified basins (4 out of 7 variants) or *t*_*R3*_ indicating slow jumps between conformational basins (3 out of 7 variants). Three variants had their middle contribution at *t*_*R2*_ arising from domain reorientations. Truncated PSG variants exhibited increased heterogeneity in dynamics (Figure 3C) although the major or middle contributions to dynamics appear at *t*_*R1*_ or *t*_*R3*_.

### Discrete Molecular Dynamics Simulations of the PSG Core

To map the conformational energy landscape of the truncated PSG supramodule, we performed replica exchange DMD simulations using 18 replicas running at neighboring temperatures with a cumulative total simulation time of 11.9 μs. To avoid bias, we chose an extended starting conformation with PDZ3 not in contact with SH3-GuK (Figure 4 – figure supplement 1). The probability density function of the radius of gyration (*R*_*g*_) shows that PDZ3 did not linger in extended conformations, which were rarely sampled (Figure 4A & Figure 4 – figure supplement 1). Instead, the PDZ3 primarily adopted a docked medium conformation (α) with a mean *R*_*g*_ of 27.6 Å along with a more compact conformation (β) with mean *R*_*g*_ of 23.4 Å. Representative models from these 3 populations (extended, medium, and compact) reveal a diverse ensemble of conformations with PDZ3 sampling both SH3 and GuK as well as undocked conformations (Figure 4B).

**Figure 4.**
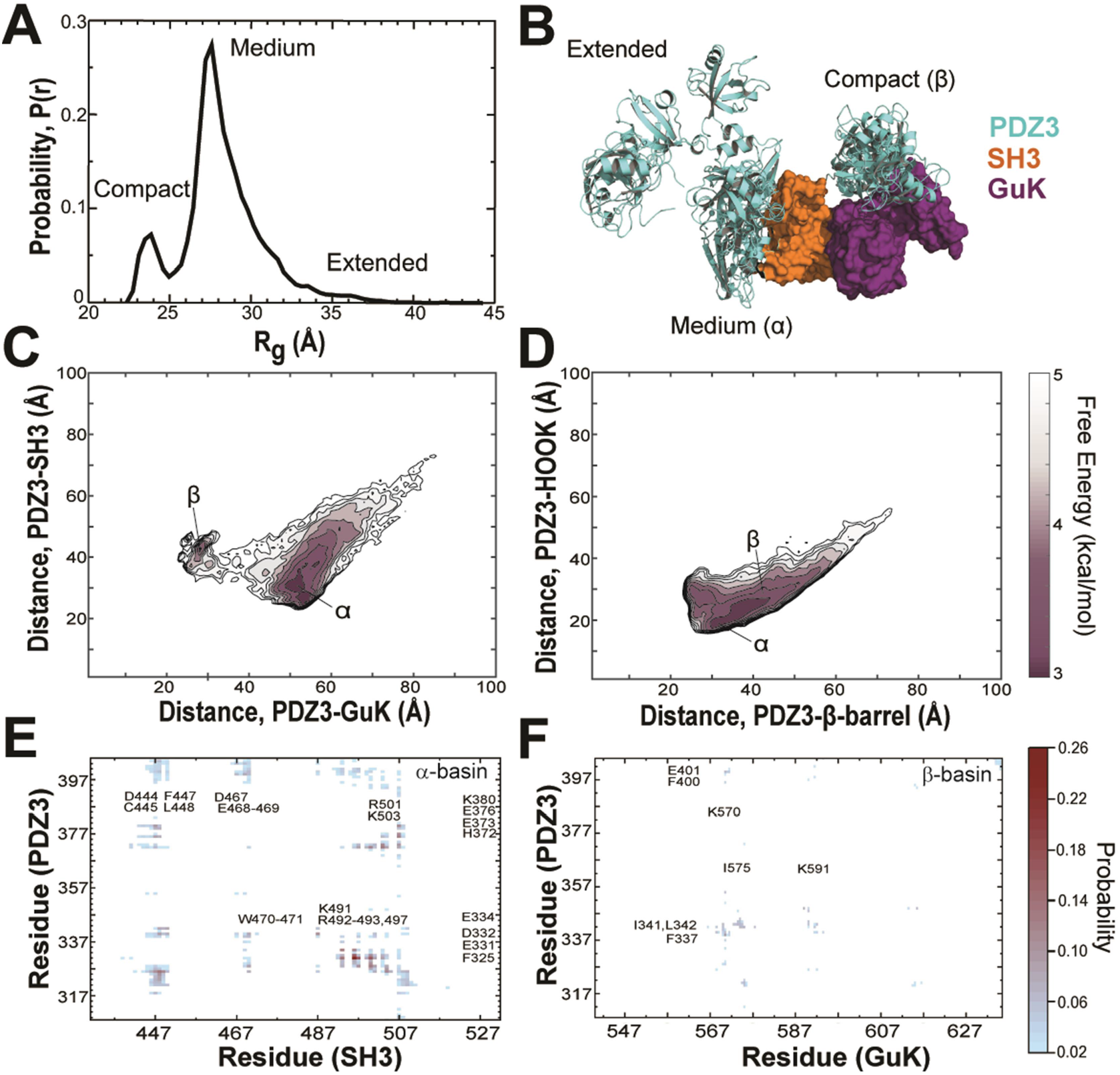
Discrete Molecular Dynamics of the PSG Supramodule from PSD-95. **A)** Probability distribution of the radius of gyration (*R*_*g*_) of the PSG during replica-exchange DMD simulations totaling 11.9 μs. **B)** Representative conformations from the 3 basins apparent in the *R*_*g*_ distribution. The conformations and their respective population fractions in the highly sampled α-basin and less frequently sampled β-basin are provided in Figure 4 - figure supplement 3). **C)** 2D free energy landscape of the relative distance between centers of mass (COM) for PDZ3 and GuK (x-axis) or SH3 β-barrel (y-axis). **D)** 2D free energy landscape of the relative distance between COM for PDZ3 and either the SH3 β-barrel (x-axis) or the SH3 HOOK insertion (y-axis). **E)** Probability of pairwise residue contacts between PDZ3 and SH3, which define the α-basin. Residues from PDZ3 are on the y-axis while residues from SH3 are on the x-axis. **F)** Probability of pairwise residue contacts between PDZ3 and GuK, which define the β-basin. Residues from PDZ3 are on the y-axis while residues from GuK are on the x-axis. The associated color bar indicates the normalized probability of the individual pairwise interactions.

To represent the cumulative association of PDZ3 with SH3 and GuK, we plotted the distance between centers of mass (COM) for PDZ3 and SH3 against the distance between COMs for PDZ3 and GuK (Figure 4C). This 2D free energy profile depicts a broad low-energy basin with PDZ3 closer to SH3 (Figure 4C, α-basin). This ensemble corresponds to the predominant population in the R_g_ distribution. Within the α-basin, PDZ3 localized to the HOOK insertion rather than the canonical SH3 domain (Figure 4D). PDZ3 also samples a shallower basin closer to GuK, which is separated by an energy barrier of ∼2.0 kcal/mol (Figure 4C, β-basin). This population corresponds to the compact configuration in the R_g_ distribution. In addition, PDZ3 samples a range of fully extended conformations with longer distances to both SH3 and GuK (Figure 4C).

Examination of structures within the α-basin reveals a broad ensemble of conformations with PDZ3 able to reorient around the HOOK helix and occasionally sample the SH3 RT loop ^33^ (Figure 4 - figure supplement 3). This suggests a fuzzy and dynamic interaction. Examination of α-basin pairwise contacts reveals that basic residues within HOOK interact with negatively charged residues in PDZ3 and are further stabilized by surface-exposed aromatic and hydrophobic residues (Figure 4E). Negatively charged residues in α3 and the β2-β3 loop of PDZ3 keep the peptide-binding face oriented towards SH3. Steric clashes would preclude PDZ3 from ligands binding while in some conformations within the α-basin (Figure 4 - figure supplement 3). PDZ3 interacts directly with SH3 rather than having this interaction mediated by the linker as had been proposed ^20^. Instead, the PDZ3-SH3 linker interacts with the SH3 domain, which helps retain PDZ3 in the α-basin and prevents more extended conformations. Hydrophobic and electrostatic interactions between the PDZ3-proximal linker (F400, E401, K403 and I404) and SH3 are among the top 50 pairwise residue contacts.

Examination of the β-basin ensemble revealed a more well-defined interaction with PDZ3 binding near the interface of the nucleoside monophosphate (NMP) binding and CORE subdomains of GuK (Figure 4B). The interaction does not occlude the canonical peptide binding sites in GuK or PDZ3 (Figure 4 - figure supplement 3). Analysis of the β-basin pairwise contacts revealed a well-defined binding site stabilized by hydrophobic and hydrogen bonding interactions between uncharged polar residues (Figure 4F) unlike the highly charged interface in the α-basin. Interestingly, the PDZ3-SH3 linker also makes significant pairwise contacts with GuK in this basin. Unexpectedly, the HOOK insertion formed an extended α-helix between residues 491 and 508 in DMD simulations. This is longer than observed in crystal structures ^12,30^ (Figure 4 - figure supplement 4). Interestingly, PDZ3 interactions appear to stabilize the helical extension as is visible in the representative basin models (Figure 4 - figure supplement 3).

### Disulfide mapping the interdomain interfaces

To confirm the PDZ3 interactions with SH3 and GuK, we engineered cysteine residues based on the contact frequency maps and measured the extent and rate of disulfide bond (DS) formation (Figure 5). DS formation depends on distance and orientation between cysteines but also contact frequency for dynamic interactions. Thus, the kinetics of bond formation report on structural proximity ^34^. We made peripheral cysteine substitutions so as to not disrupt the primary contacts, which are shown using the structure with the lowest root mean square deviation (RMSD) to each basin ensemble. The predominant interface in the α-basin was PDZ3 binding to the HOOK. We probed this basin with the residue pair N326C-A504C, which are only 5.6 Å apart in the α-basin representative (Figure 5A). The predominant interface in the β-basin had PDZ3 binding to GuK. We probed this basin with the residue pair G344C-N633C, which are 5.7 Å apart in the β-basin representative (Figure 5B).

**Figure 5.**
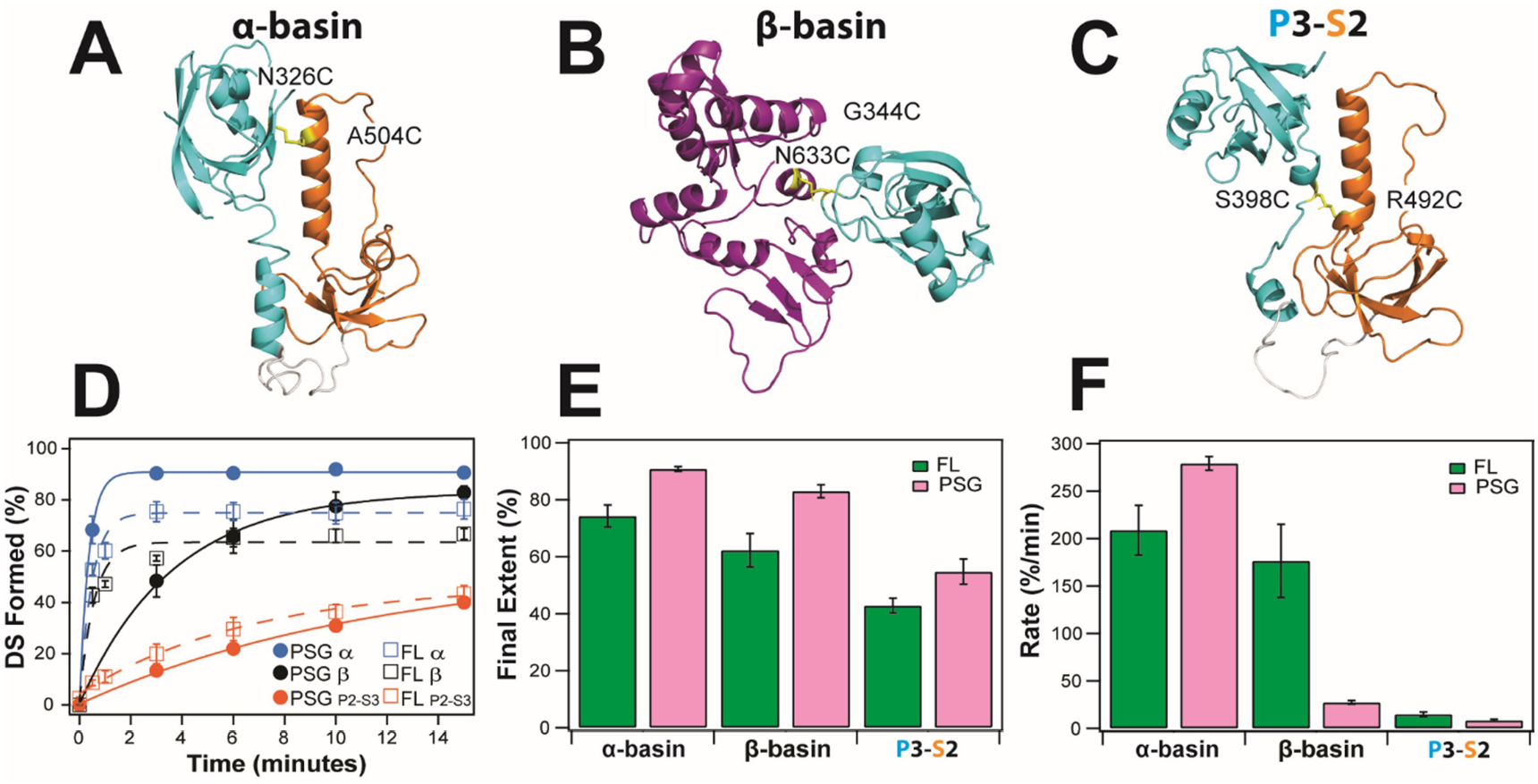
Disulfide Mapping of the Contact Interfaces Identified from DMD Simulations. Cartoon representations showing PDZ3 (cyan), SH3 (orange) and GuK (purple). **A)** Target model from the α-basin. N326C-A504C disulfide in yellow. **B)** Target model from the β-basin. G344C-N633C disulfide in yellow. **C)** Predicted disulfide binding contacts involving FRET variant P3-S2. S398C-R492C disulfide in yellow. **D)** Kinetics of disulfide bond formation as measured using non-reducing SDS-PAGE (Figure 5 - figure supplement 1) showing the α-basin (blue), the β-basin (black) and variant P3-S2 (orange). Data for full-length PSD-95 are shown as circles with fits as solid lines while data for the PSG truncations is shown as open circles with dashed lines. **E)** Final extent of disulfide bond formation from single exponential fits to the kinetic data for full-length PSD-95 (green) and the PSG truncation (pink). **F)** Kinetic rate of disulfide bond formation from single exponential fits to the kinetic data. Error bars represent the standard deviation from three replicate measurements.

Increased electrophoretic mobility indicated that DS formation was occurring for all samples (Figure 5 - figure supplement 1). The data were fit to an exponential function to determine the rate and final extent of DS formation (Figure 5D). The α-basin variant showed slightly more DS formation than the β-basin variant in full-length PSD-95 but the rates of DS formation were similar (Figure 5E & F). To probe the effects of truncation, we measured DS formation in the PSG truncation. Interestingly, the truncation had opposite effects on the kinetics of DS formation for the two variants. The rate of DS formation for the α-basin variant increased by ∼30% while rate of DS formation for the β-basin variant decreased by six-fold.

We also screened the DMD simulations for FRET variants capable of DS formation, which predicted that variant P3-S2 occasionally sampled close distances in the α-basin (Figure 5C). As predicted, variant P3-S2 formed DS albeit to a lesser extent and much more slowly than either designed variant. The extent of DS formation was slightly higher in the truncation, but the rates were similar. Thus, DS formation for all three variants was higher in the truncation. Our DS analysis confirms that the contact interfaces from DMD are sampled in full-length and truncated PSD-95. We also observe significant kinetic differences when PSD-95 is truncated in agreement with FRET studies.

### Structural Modeling with Experimental FRET Restraints

To describe the conformational ensemble associated with each limiting state from seTCSPC, we used the FRET distances as restraints in structural modeling. We simulated the accessible volume (AV) for both dyes at each labeling site ^35^ (Supplementary file 4A), which we used as distance loci for rigid body docking. For each state, we generated 33000 conformations, which were each scored for agreement between the AV model distances and the FRET distances. For state A, the best-fit models showed some divergence with PDZ3 near the HOOK insertion but also extended without interdomain contacts (Figure 6A). The best-fit models for state B are more tightly clustered and position PDZ3 near the NMP subdomain of GuK (Figure 6B).

**Figure 6.**
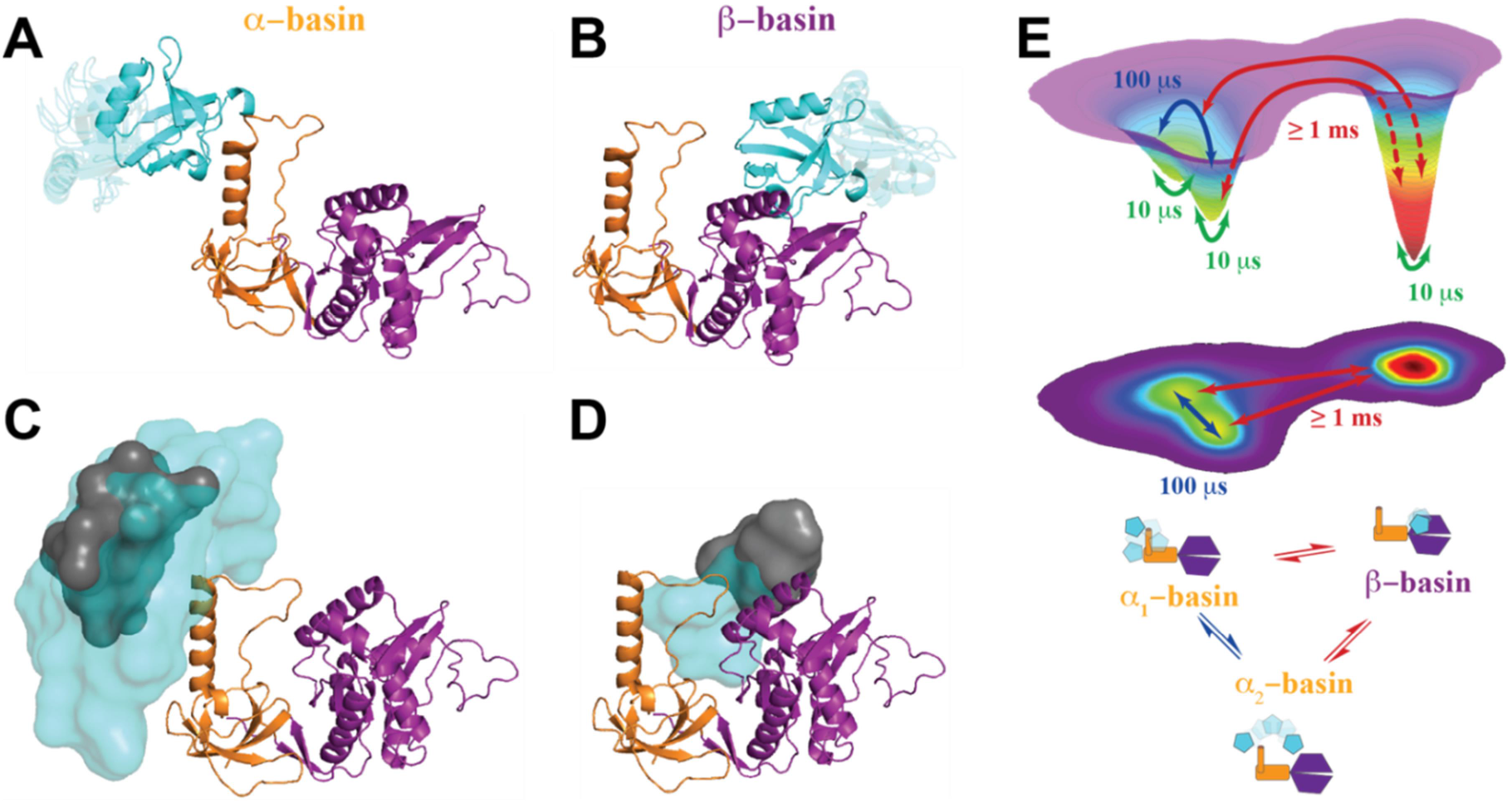
Modeling the Supertertiary Structural Ensemble with Experimental FRET Restraints. **A-B)** Cartoon representations of the best fit models from rigid body docking of PDZ3 based on the FRET distances from seTCSPC (PDZ3, cyan; SH3; orange; GuK, purple). **A**) Best fit model for full-length PSD-95 in limiting state A. (**B**) Best fit model for limiting state B. **C-D)** Heterogeneity of the conformational state ensembles based on classification of structures from DMD trajectories into state A (**C**) or state B (**D**). The grey surfaces represent 95% confidence intervals for localization of the PDZ3 center of mass based on screening with limiting state distances from all 12 full-length variants. Confidence intervals are based on the F-test for the ratio of χ^2^_r_ for all docking structures relative to the χ^2^_r_ of the top-ranked structure with 9 free parameters (number of distances used for docking of PDZ3). The grey surfaces represent only the uncertainty in distances from the global fit but do not capture the full heterogeneity of each basin. The cyan surfaces represent conformational space accessible to the PDZ3 center of mass from screening DMD with thresholds from FNR reanalysis of seTCSPC data (Figure 6 – figure supplement 1 & 2, Supplementary file 4B). **E)** Energy landscape for the conformational ensemble of the PSG supramodule within PSD-95. Transitions and their associated timescales taken from fFCS indicated by colored arrows. The fastest transitions are associated with local motions, which were not resolved by FRET but are inferred from DMD simulations and fFCS. The energy landscape surface was generated by rescaling the surface that resulted from principal component analysis (Figure 6 – figure supplement 3) such that the integrated volumes of basins α and β matched the population fractions for states A and B, respectively, from TCSPC analysis. Basins α_1_ and α_2_ correspond to the two sub-basins separated by a small shoulder along principal component 2 (Figure 6 – figure supplement 3).

To independently corroborate the docking models, we calculated the AV for all snapshots structures from the DMD trajectory. For each structure, we plotted the inter-AV distance (⟨*R*_*DA*_⟩_*AV*_) against the distance between the COM of labeled domains (Figure 6 – figure supplement 1). To denote the limiting states, we overlay orange and purple vertical lines for state A and B respectively. The limiting states A and B generally fall within the associated α- and β-basins. However, it was also clear that for some variants, (e.g. P2-G6) the vertical line for state B agrees with both DMD basins. Similarly, the PDZ3-SH3 variants may not fully capture the underlying population distribution. This apparent discrepancy rises from what we call FRET degeneracy in which a single distance captures an underlying heterogenous distribution. To resolve the FRET degeneracy, we introduce the FRET Network Robustness (FNR) analysis (Figure 6 – figure supplement 2). We systematically refit the seTCSPC data using different numbers and combinations of variants from the FRET network. We randomly selected sub-samples with as few as 5 variants while including each variant in at least 3 subsets. The distance deviation increased with fewer variants, but the FNR distributions remained centered on the distance from the full global fit (Figure 6 – figure supplement 2). When more than 7 variants were used, the FNR deviation was less than 10% regardless of which variants were included in the subset. The standard deviation of the FNR distributions (along with AV simulations) captures the heterogeneity introduced by FRET degeneracy, which can be used to impose new bounds on structural heterogeneity (Supplementary file 4B).

Using these bounds, we screened the 20871 snapshot structures from DMD simulations to classify all models that fall within these new bounds for the α- and β-basins. Displaying the PDZ3 COM for each compatible model as a cyan sphere emphasizes the fuzziness of the basins as captured by the underlying heterogeneous FRET distributions from each variant. The broad ensemble for the α-basin is quantified by the variance of the FNR distributions (Figure 6 – figure supplement 2) but arises from the fuzziness of interdomain interactions between PDZ3 and SH3 (Figure 6C). In contrast, the β-basin is not fuzzy (Figure 6D). For comparison, we overlay the COM of screened DMD snapshots that fall within the 95% confidence interval excluding contributions from FRET degeneracy (Figure 6C and 6D; grey spheres).

We used this information to construct a conformational landscape consistent with DMD and FRET (Figure 6E). We performed principal component analysis and projected the basins along the first two principal components (PC1 and PC2) using the COM distances and the simulated ⟨*R*_*DA*_⟩ between PDZ3 and SH3-GuK, which were rescaled such that the integrated volume of each basin was equivalent to its population fraction from seTCSPC (Figure 6 – figure supplement 2). PC1 separated the α- and β-basins mostly by interdomain displacement, while PC2 describes the α-basin FRET degeneracy due to domain reorientations within a single basin. Hence, we expand the energy landscape to include degenerate α_1_- and α_2_ basins and link the fFCS relaxation rates directly to the barrier heights for transitions between basins.

### Effect of PDZ3 Native Context on Interactions with Neuroligin

Our PSG models include conformations that could impact ligand binding to PDZ3. To test this, we examined the interaction with neuroligin 1a, a key synaptic adhesion protein that interacts with PDZ3 (Figure 7A) ^24,25^. To compare binding between the truncated PDZ3 and full-length PSD-95, we used a 10 residue C-terminal neuroligin peptide (NL10), which bound truncated PDZ3 with an equilibrium dissociation constant (*K*_*D*_) of 10 µM ^26^. We N-terminally-labeled the peptide with fluorescein and measured the fluorescence anisotropy.

**Figure 7.**
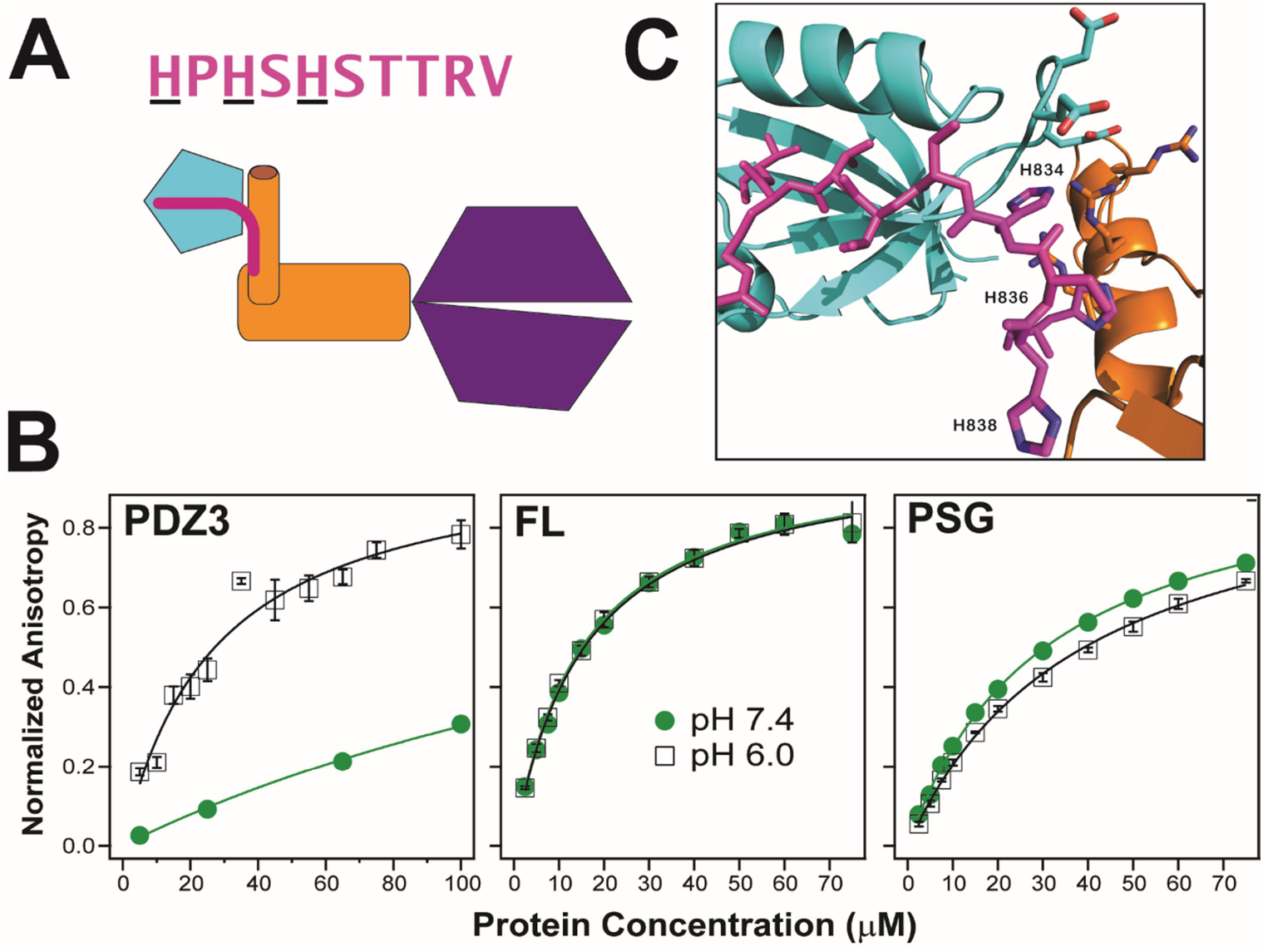
Effect of Supertertiary Environment on Neuroligin Binding. **A)** Cartoon schematic illustrating interaction between PSD-95 and neuroligin (pink). The sequence of the neuroligin C-terminal peptide used in binding experiments is shown with pH-sensitive histidines underlined. **B)** Representative binding isotherms for the neuroligin peptide binding to truncated PDZ3 (left), full-length PSD-95 (middle) and truncated PSG (right). Shown are data at pH 7.4 (green circles) and pH 6.0 (open squares). Anisotropy values were normalized relative to the final anisotropy taken from non-linear least squares fits (lines). Error bars represent the standard deviation from three replicate measurements. **C)** Representative conformation from docking of the neuroligin peptide (pink) to PSG in the α–basin from DMD (Figure S18). The peptide C-terminus interacts with the GLGF motif in PDZ3 (cyan). Charged residue interactions between PDZ3 and SH3 (orange) prevent electrostatic repulsion of histidines that otherwise weakens peptide binding at neutral pH.

We obtained a *K*_*D*_ of 25 ± 4 µM at pH 6, which is in good agreement given our higher ionic strength. Surprisingly, we were unable to reach saturation when we repeated this measurement at pH 7.4 indicating that the *K*_*D*_ increased to over 200 µM (Figure 7B). Thus, neuroligin binding showed strong pH dependence and bound to PDZ3 poorly at physiological pH. In contrast to truncated PDZ3, full length PSD-95 gave a *K*_*D*_ of 15 ± 1 µM at pH 6 and pH 7.4 suggesting the NL10 interaction is enhanced by the supertertiary environment of PDZ3. Next, we examined neuroligin binding to the PSG truncation. We measured a *K*_*D*_ of 39 ± 1 µM at pH 6 suggesting that binding affinity is somewhat reduced in the PSG relative to full length or even the PDZ3 truncation. The PSG showed slightly higher binding affinity at pH 7.4 with a K_D_ of 31 ± 2 µM in stark contrast to the reduced binding affinity of the truncated PDZ3. Thus, effects of truncation on supertertiary structure and dynamics impact NL10 binding.

To understand this phenomenon, we performed docking of NL10 to truncated PDZ3 at pH 6. We identified electrostatic interactions between protonated histidines in NL10 and acidic residues in the PDZ3 β2-β3 loop (Figure 7 – figure supplement 1). At pH 7.4, unprotonated histidines are incapable of interacting, which explains the pH sensitivity. The resulting desolvation penalty of the unpaired acidic residues would reduce the binding affinity. Some α-basin conformations block peptide binding due to steric clashes. However, docking of NL10 to α-basin structures identified multiple PSG conformations where the acidic residues in PDZ3 interacted with basic residues in HOOK without steric clashes for NL10 (Figure 7C). Salt-bridges have a smaller desolvation penalty than unsatisfied charges, which explains why the peptide binds full-length PSD-95 (and PSG) better than truncated PDZ3 at pH7.4. Thus, the supertertiary context of PDZ3 enables recognition of a critical physiological ligand.

## DISCUSSON

PSD-95 is a scaffold protein at excitatory synapses that forms a crucial link between neurotransmitter release and detection pathways ^6^. PSD-95 brings together different binding partners from oligomeric transmembrane proteins to soluble enzymes. Proteomic analysis of PSD-95 complexes purified from mouse brain identified 301 different proteins ^36^. In many cases, the binding partners are larger than the scaffolding domains with which they interact. Kinetic analysis showed that higher-order interactions, between proteins bound to PSD-95, plays a role in scaffolding activity ^37^. The assembly of multi-protein complexes poses steric challenges. Thus, the dynamic supertertiary structure must play a role in accommodating multiple partners.

Here we used a combination of experimental and computational methods to describe the supertertiary structure and dynamics of the conserved PSG supramodule within full-length PSD-95. Multiparameter fluorescence analysis revealed a complex and dynamic conformational landscape. DMD and modeling based on FRET distances were in excellent agreement on the location of PDZ3 within two non-overlapping basins. Both DMD and FRET agree that the α-basin was degenerate due to the fuzzy interface allowing PDZ3 reorientation. The fFCS was crucial for assigning dynamic timescales to the conformational transitions, which we summarized as an energy landscape (Figure 6E). Within each basin, each domain undergoes rapid local motions. Reorientation of PDZ3 and sampling of extended states is slower while basin exchange take place on timescales approaching seconds as captured by smTIRF.

Our models were in excellent agreement with SAXS and NMR experiments, which identified multiple conformations with PDZ3 predominantly localized near the HOOK insertion and suggested involvement of residues around the PDZ3 peptide binding pocket ^20^. However, Rosetta modeling suggested the PDZ3-SH3 linker bridged the interaction while we found the PDZ3-SH3 interaction was direct. The linker interacts with SH3 and GuK to help retain PDZ3. This linker is almost 100% conserved in PSD-95 homologues from humans to drosophila, which would be unusual unless involved in protein interactions since disordered linkers usually show reduced sequence conservation ^38^. Mutations in the PDZ3-SH3 linker could explain the reduced interactions seen in NMR on re-engineered PSG constructs ^39^. Neither basin relied on canonical ligand binding modes for the primary interaction as suggested by comparative patch analysis ^21^.

The crystal structure of the PSG supramodule from ZO-1, a MAGuK protein found at intercellular tight junctions, revealed a lack of interdomain interactions ^40^. The PDZ3-SH3 linker in ZO-1 is much shorter than PSD-95, which may prevent access to these binding sites. The ZO-1 HOOK insertion also lacks sequence similarity to PSD-95 ^40^. In contrast, the basic residues in HOOK and acidic residues in PDZ3 are conserved among synaptic MAGuK homologs PSD-93, SAP97, and SAP102 as are most pairwise contacts in the β-basin (e.g., F339-Q594, L342-Q603). This explains why the supertertiary landscape is conserved in these homologues as suggested by previous smTIRF measurements ^22^.

The importance of supertertiary interactions on scaffolding activity is emphasized by the enhanced binding of neuroligin to full-length PSD-95. Other PDZ3 ligands (e.g., CRIPT, stargazin and synGAP) contain lysines and arginines near the canonical PDZ motif while neuroligin contains histidines, which explains why NL10 has pH sensitivity. In the fuzzy α-basin, interactions with positive charges in the HOOK satisfy negatively charged residues in PDZ3, which facilitates uncharged peptide binding. Additionally, fuzziness increases the favorability of the α-basin by allowing PDZ3 to reorient to satisfy multiple interactions. These supertertiary interactions also affect other ligands. Measurements of CRIPT binding to PDZ3 suggested that PDZ3 adopted two interconverting conformational states in the PSG with different kinetics but the same equilibrium affinity ^41^. Thus, the supertertiary context of PDZ3 is necessary to overcome the repulsive interactions that prevent neuroligin binding to the truncated domain.

## METHODS

### Protein expression and purification

The full-length PSD-95 from *Rattus norvegicus*, the PSG truncation (residues 303-274) and truncated PDZ3 (303-415) were expressed in the Rosetta 2 strain of *E. coli* (EMD Millipore) and purified by a combination of Ni-affinity, ion exchange and size exclusion chromatography as previously described ^14^. For each construct, we prepared a series of labeling variants using standard site directed mutagenesis and confirmed with DNA sequencing (Table 1). For smTIRF, proteins were labeled Alexa Fluor 555 C_2_ maleimide and Alexa Fluor 647 C_2_ maleimide at an equimolar ratio. For MFD, proteins were first labeled with a 1:2 ratio of Alexa 488 C_5_ maleimide for 1 hour at 4°C followed by addition of a 5:1 molar ratio of Alexa Fluor 647 C_2_ maleimide, which was reacted overnight at 4°C. Unconjugated dye was removed by desalting with Sephadex G50 (GE Healthcare) followed by dialysis.

### Single-Molecule Total Internal Reflection Fluorescence (TIRF)

Fluorescently labeled PSD-95 was encapsulated in 100 nm liposomes supplemented with 0.1% biotinylated phosphoethanolamine (Avanti Polar Lipids, Alabaster, AL). Unencapsulated protein was removed by desalting on Sepharose CL-4B (GE Healthcare). Liposomes were attached via streptavidin to a quartz slide passivated with biotinylated-BSA. FRET data was collected at 10 frames/second using an Andor iXon EMCCD camera (Andor Technologies, Belfast, UK). All smFRET measurement were performed in 50 mm Tris 150 mM NaCl pH 7.5 supplemented with 5 mM cycooctatetraene, 0.5% w/v glucose, 7.5 units/mL glucose oxidase and 100 units/mL catalase. Alternating illumination using 640 and 532 nm lasers allowed for the identification of molecules containing one donor and one acceptor. Microscopy data was analyzed using home written scripts in MATLAB to correlate donor and acceptor images, extract single-molecule intensity time traces from the pixel intensity and calculate FRET efficiency with corrections for background and crosstalk. Automated per molecule gamma normalization based upon acceptor photobleaching events was used to correct for distortions in the measured intensities between the donor and acceptor channels ^42^.

### Multiparameter Fluorescence Detection

Two experimental setups were used for confocal measurements to obtain MFD data (Supplementary file 1A & B). For the Clemson University setup, freely diffusing molecules were excited as they passed through the focal volume of a 60X, 1.2 NA collar (0.17) corrected Olympus objective with diode lasers at 485 nm and 640 nm (PicoQuant, Germany) operating at 40 MHz with 25 ns interleaved time. The power at the objective was 80 µW at 485 nm and 32 µW at 640 nm. Emitted photons were collected through the same objective and spatially filtered through a 70 µm pinhole to limit the effective confocal detection volume. For the setup located at Heinrich Heine University (HHU), molecules were excited with diode lasers at 485 nm and 640 nm (PicoQuant, Germany) operating at 64 MHz (15.6 ns interleaved time). The power at the objective was set to 60 µW at 485 nm and 10 µW at 640 nm. The location at which each sample was measured is indicated in Supplementary file 1A.

Emission was separated into parallel and perpendicular polarization components at two different spectral windows using band pass filters, ET525/50 and ET720/150, for donor and acceptor, respectively (Chroma Technology Co.). In total, four photon-detectors are used at the Clemson setup—two for donor (PMA Hybrid model 40 PicoQuant, Germany) and two for acceptor (PMA Hybrid model 50, PicoQuant, Germany). To ensure temporal data registration of the 4 synchronized input channels, we used a HydraHarp 400 TCSPC module (PicoQuant, Germany) in Time-Tagged Time-Resolved mode. For the HHU setup, eight detection channels are used—four for green (τ-SPAD, PicoQuant, Germany) and four for red APD SPCM-AQR-14, Perkin Elmer, Germany) data registration synchronization was achieved with HydraHarp 400 TCSPC module (PicoQuant, Germany) in Time-Tagged Time-Resolved mode.

Both setups utilized Pulsed Interleaved Excitation (PIE) to alternately excite donor and acceptor fluorophores directly^43^. Emission was separated into parallel and perpendicular polarization components at two different spectral windows using band pass filters as described in supplementary methods. Labeled samples were diluted in 50 mM sodium phosphate, pH 7.5, 150 mM NaCl, 40 µM TROLOX, which had been charcoal filtered to remove residual impurities. We used pM concentrations such that we observed ∼1 molecule per second in the confocal volume. Samples were measured in NUNC chambers (Lab-Tek, Thermo Scientific, Germany) that were pre-coated with a solution of 0.01% Tween 20 (Thermo Scientific) in water for 30 min to minimize surface adsorption. We obtained the instrument response function (IRF) using water. Protein-free buffer was used for background subtraction. Calibration experiments and data collection were as previously reported ^44^. Burst selection was performed using all-photon, inter-photon arrival time traces to identify single molecules. Burst selection criteria were set such that each burst contained a minimum of 40 photons summed amongst all detection channels, with an inter-photon arrival time cutoff set to the mean minus one standard deviation, calculated across the entire time trace. The donor fluorescence lifetime and the intensity-based FRET efficiency were calculated for each burst using a maximum-likelihood estimation algorithm ^23,45^. To ensure both fluorophores were present in each select burst used in MFD histograms, we used cutoff values for 1) the difference between observed burst duration in green and red channels under direct excitation of the corresponding fluorophores (|T_GG_-T_RR_|<1 ms) and 2) the observed FRET stoichiometry (0.3<S_PIE_<0.7) ^23^.

### Subensemble Time-Correlated Single-Photon Counting (seTCSPC) Analysis

Analysis of donor fluorescence decays in conditions for FRET to determine limiting states was performed on photon data corresponding to bursts shown in Figure 1 – figure supplement 3. These fits and raw data are shown in Figure 1 – figure supplement 2 and Figure 2 – figure supplement 2 for full-length and truncated PSG core, respectively. Data for fits was generated by loading burstwise data and IRF data corresponding to the green detection channels into the FitMachine analysis software using 4096 TAC bins and saving the resulting fluorescence decay and IRF histograms. Fitting was performed in the ChiSurf analysis software, which allows users to set individual parameters to be shared globally and optimized amongst subsets of curves. Fit models were multi-exponential decays described by:

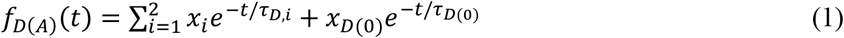

where *x*_*i*_ and 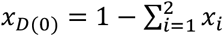 are species fractions of exponential decay terms for FRET-sensitized and DO/no-FRET emission, and *τ*_*D,i*_ and *τ*_*D*(0)_ are the corresponding donor fluorescence decay lifetimes. The instrument response function (IRF) is deconvolved and decays are related to states with underlying Gaussian-distributed interdye distances as described ^23,46^. Prior to global fitting, donor-only (DO) fluorescence lifetimes and fractions were determined by fitting each variant with the corresponding DO curves. These DO curves are shown in Figure 1 – figure supplement 2 and Figure 2 – figure supplement 2, and the DO lifetimes and fractions are provided in Supplementary file 1D. Global fits for the full-length variants were performed by loading all decay data simultaneously along with the corresponding IRF data. DO fluorescence lifetimes, DO fractions, and instrumental parameters were fixed for each sample based on DO fits, and two distances corresponding to donor fluorescence lifetimes and fractions were used for each FRET variant, with population fractions set as global parameters and distances optimized individually. The fit parameters for this analysis are listed in Supplementary file 1D. Where *τ*_*D,xi*_ and *τ*_*D,fi*_ are used, *x* indicates the species mean fluorescence lifetime used in seTCSPC fits and *f* indicates the fluorescence-weighted average lifetime for the state used in computing FRET-lines for MFD histograms. Conversion to distances was performed as we have done previously, using a standard Förster radius of 52 Å for the Alexa 488 and Alexa 647 dye pair (2). The Förster radius was corrected individually for each sample based on the corresponding mean dye orientation factor, ⟨*κ*^2^⟩, given in Supplementary file 1D. The expression for ⟨*κ*^2^⟩ and relation to the Förster radius has been detailed previously ^47^. The values of ⟨*κ*^2^⟩ were estimated using the Kappa2-Distribution tool in the Chisurf software package. The values of steady-state anisotropy used for these estimations (r_D|D,∞_, r_A|A,∞_ for donor and acceptor under direct excitation, respectively) were calculated from MFD histograms and are provided in Supplementary file 1D. Interdye distances from the seTCSPC fitting can be found in Supplementary file 1E. This procedure was repeated for the truncated PSG variants.

### Photon Distribution Analysis

PDA analysis accounts for shot noise, background signal contributions, and binning effects to construct accurate fluorescence parameter histograms ^31,48^. PDA analysis was performed using the Seidel Software Tatiana 4.8 software. For this analysis, burstwise photon data was re-binned into time windows of varying lengths: 1 ms, 2 ms, and 3 ms. Shorter time windows corresponding to the ends of bursts partway through a time window were discarded. Burstwise FRET efficiency was recalculated from this re-binned data. The resulting histograms were fit globally for each FRET variant to ensure the resulting model is consistent across different time windows sizes. The model used included two static populations corresponding to the limiting state distances with fixed widths (7% of R_D(A)_) as well as a dynamic population corresponding to exchange between these states. Fixed-width states were included to account for apparently static states corresponding either to conformationally static states or dynamic processes not reflected by changes in the FRET efficiency of any given variant. The reaction rates, *k*_AB_ and *k*_BA_ as well as the population fractions, A_A_, A_B_, and A_Dyn._ were allowed to vary. This model can be summarized as (*A* ↔ *A*′) ↔ (*B* ↔ *B*′). The mean relaxation time is given by 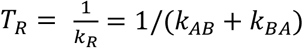. Correction parameters for PDA FRET efficiency histograms were the same as for multiparameter histograms (Supplementary file 1A & B).

### Filtered Fluorescence Correlation Spectroscopy (fFCS)

For modeling the fFCS we use the following functions.

Species Auto-Correlation Function:

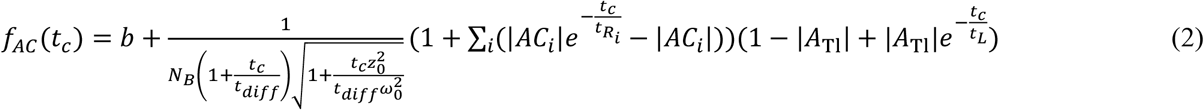

Species Cross-Correlation Function:

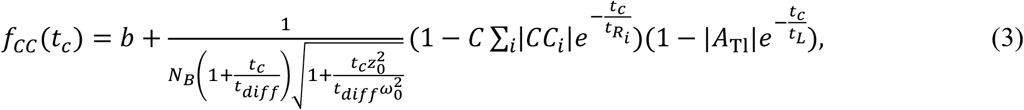

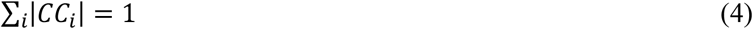

Here, *b* is the baseline value for the correlation function, corresponding to no correlation, *N*_*B*_ is the average number of bright molecules in the confocal volume, *t*_*c*_ is the correlation time, *t*_*diff*_ is the average diffusion time through the confocal volume, *s* = *z*_0_/*ω*_0_ is a geometric factor of the confocal volume as the ratio of widths parallel and perpendicular to the light path of the excitation laser through the sample, *AC*_*i*_ and *CC*_*i*_ are auto-correlation and cross-correlation decay amplitudes, 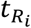 are the associated decay times, *C* is a normalization factor for each individual cross-correlation curve, and *A*_Tl_ and *t*_*L*_ are additional decay components to account for long-timescale processes not associated with dynamics that occur below the diffusion time.

### Structure generation and system setup

The PDZ3 and SH3-GuK domains of the PSG core were reconstructed using homology modeling of I-TASSER (Iterative Threading ASSEmbly Refinement) hierarchical approach using multiple threading alignments and iterative structural assembly simulations. The I-TASSER method involves four consecutive stages: (a) template identification or threading by LOMETS, (b) structure assembly by replica-exchange Monte Carlo simulations, (c) model selection and structure refinement using REMO and FG-MD, and (d) structure-based function annotation using COFACTOR ^49 50^. The individual PDZ3, SH3-GuK domains were positioned randomly and our in-house loop reconstruction program ‘medusa-loop’ was utilized to connect the individual domains via flexible linkers to reconstruct the PSG core (Figure 4 – figure supplement 1). The backbone of the GuK domain was kept static while the SH3 and PDZ3 domains were kept flexible to allow adequate sampling of the conformational landscape. Gō constraint was imposed on the intra-PDZ3 (residues 308-405), intra-SH3 (residues 431-531) domains and the β strand F of GuK (residues 710-724) that folds back to form an antiparallel β sheet with β strand E of the SH3 domain. The implementation of Gō-like constraints, based on inter-residue contacts, can provide a reliable model consistent with the experimentally-established native structure of PDZ3 and SH3-GuK.

### Replica exchange DMD simulations

Replica exchange DMD (rxDMD) simulations were performed with 18 neighboring replicas in the temperature range of 275-360 K to efficiently sample the conformational free energy landscape. In the rxDMD scheme, temperature was exchanged between replicas at periodic time intervals, according to the Metropolis criterion, to overcome any local energy barriers that may limit efficient sampling of the conformational states. Production runs followed 3000 timestep energy minimization runs, where a DMD timestep corresponded to ∼50 fs. The conditions for replica exchange were checked every 1000-time units and the frames were saved every 100-time units. Anderson’s thermostat was used to maintain temperature at 300 K in all simulations and the heat exchange factor was set to 0.1. Each replica simulation lasted for 660 ns that resulted in a total simulation time of ∼11.9 *μs*.

The thermodynamics of the PSG core was computed using the Weighted Histogram Analysis Method (WHAM) ^51,52^. The time evolution of the *R*_*g*_ for the 18 simulated replicas is provided in Figure 4 – figure supplement 2. We observed dynamic, spontaneous conformational exchange in each of the replicas (Figure 4 – figure supplement 2), highlighting the efficient sampling from exchanging temperature between replicate simulations. The 2D free energy landscape was computed as a function of the interdomain distance between PDZ3-SH3_Hook_, PDZ3-SH3_barrel_ and PDZ3-GuK domains. Using the last 400 ns of replica exchange trajectories, the density of states *g*(*E*) was calculated by combining energy histograms from all 18 replicas. The 2D potential of mean force (PMF) as a function of interdomain distance was computed as:

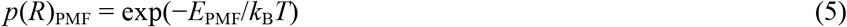

where *E*_PMF_ is the free energy integrated over the interdomain distance, *R. T* is the temperature of interest and *k*_B_ is the Boltzmann constant. The free energy contours are scaled in units of kcal/mol.

The hierarchical clustering program (www.compbio.dundee.ac.uk/downloads/oc) was utilized to group similar conformations of the PSG core. Depending on the pair-wise root-mean square distance (RMSD) matrix, the clustering algorithm iteratively combines nearby clusters. The “cluster distance” was determined based on all pairwise distances between elements of corresponding clusters. We used the mean of all values to compute the distance between two clusters, and the centroid structure of each cluster was chosen with the smallest average distance to other elements in the cluster.

The disulfide bond modeling of the interdomain contacts in PSG was performed for the two basins (α and β) in the free energy landscape. From the structural ensembles corresponding to the reaction coordinates in the residual contact frequency map, we ranked contact residue pairs and selected the top three pairs near the interface that did not involve the hydrophobic and salt-bridge interactions. The selected pairs were situated at the peripheral sites of the interdomain interface, and the corresponding residues were either polar or weakly hydrophobic in nature ^34^.

### Rigid Body Docking and Screening of DMD Structures

Rigid body docking and screening against DMD simulations were performed using the FRET Positioning and Screening (FPS) software ^27^. Input distances and uncertainties were taken from seTCSPC in Supplementary file 1E. Initial structure was generated from the representative DMD structure used throughout the main text by removing residues 399-428 corresponding to the PDZ3-SH3 linker such that PDZ3 and SH3-GuK are treated as separate rigid bodies. In simulations, PDZ3 is held static while SH3-GuK undergoes a biased random walk with FRET distance restraints imparting forces to guide the docking toward those structures satisfying the restraints. Distances are calculated for rigid bodies by sampling fluorophore accessible volumes (AV) simulated for each fluorophore labeling position. 11000 independent runs were performed from the initial structure for each limiting state distance set from global seTCSPC fitting. All resulting structures for these runs were combined into one dataset and best structure selection was performed from this combined dataset. Representative structures are identified based on minimization of 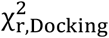 calculated between simulated distances for each structure and experimental distances. This procedure was repeated for the distances from the truncated variants. For contact state visualization, an additional restraint per successful DS pair was included for the structure with the shortest distances corresponding to each DS pair, set to 2 Å +/- 2 Å.

Screening and FRET robustness analysis were performed in the same software by loading 20871 structures from DMD simulations, simulating AVs for each labeling site in every structure, calculating inter-dye distances by random sampling of the AVs, and again using distances and uncertainties from seTCSPC for comparison. These structures were not modified and therefore included all of residues 308-724. Best representatives were determined for screening in the same way as for docking structures but using 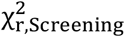. Analysis of refitted distances from subsamples of variants were analyzed using the same AV data as for screening. For each structure, interdye distances were calculated from dye AVs and structures were classified based on whether the average percent error of simulated distances for that structure relative to the distances in Supplementary file 4B were below the thresholds corresponding to either basin A, basin B, or neither. Surfaces were calculated corresponding to the PDZ3 center-of-mass positions for all structures classified into either basin, shown in Figure 6. Contours representing classifications of structures are shown in Figure 6 – figure supplement 1. Resulting structures and surfaces from these analyses were visualized in the PyMOL Molecular Graphics System, Version 2.0 (Schrödinger, LLC). Attachment sites and parameters for AV simulations for both docking and screening can be found in Supplementary file 4A.

### Disulfide Mapping in PSD-95

All samples were pre-reduced with 5 mM DTT for one hour followed by desalting using a PD10 column (GE Healthcare) into 20 mM tris pH 7.4, 150 mM NaCl. Disulfide oxidation reactions were performed at 25°C with 2 μM protein concentration for the PSG fragment or 0.5 μM for full-length PSD-95. Disulfide formation was initiated by the addition of 0.5 mM CuSO_4_ and 1.75 mM 1, 10-phenanthroline. Time points were quenched by adding 40 mM N-ethylmaleimide and 10 mM EDTA. Samples were boiled in non-reducing Laemmli sample buffer and run on 10% or 5% SDS-PAGE for PSG and full-length, respectively. All experiments were carried out in triplicate. Intensities for native and shifted bands were measured in ImageJ. Percentages of disulfide formation were calculated for each time point and corrected for the presence of higher order oligomers. Each reaction was well fit to a single exponential function to obtain the initial and final extent of disulfide formation along with the reaction rate for each mutant. Replicates were analyzed separately to obtain the average and standard error of measurement (SEM) as well as to estimate the error in the fitted parameters.

### Neuroligin Peptide Binding

Binding experiments used protein constructs described above with a 10-residue synthetic peptide corresponding to the C-terminal 10 residues from rat neuroligin 1a (HPHSHSTTRV), which was synthesized with an N-terminal fluorescein label (5-FAM, GenScript USA Inc. Piscataway, NJ). Fluorescence polarization measurements were carried out in black 96-well plates measured on a Wallac Victor 2 Plate Reader (Perkin Elmer). For all measurements peptide was held at concentrations of 50 nm (or 100nM) while the protein concentrations ranged from 1-100μM. Measurements at pH 7.4 were made in 20mM Tris, 150mM NaCl, 1mM DTT and 1mM EDTA. Measurements at pH 6.0, were made in 20mM MES, 150mM NaCl, 1mM DTT and 1mM EDTA. Four readings were taken for each well, then averaged. All experiments were carried out in triplicate and fitted with the Hill Equation for a single site binding site model.

## Code Availability

MFD is made available at http://www.mpc.hhu.de/en/software. DMD simulation engine is available at http://www.moleculesinaction.com.

## Acknowledgements

This work was supported by NIH (MH081923 to MEB, 1P20GM130451 to HS, and GM119691 to FD), NSF (CAREER MCB 1749778 to HS and CAREER CBET-1553945 to FD); and ERC (AdG2014 hybridFRET (# 671208) and HHU Connect to GH, HS, and CS. We thank Markus Seeliger (NIH S10OD028478) and Aziz Rangwala for technical assistance with peptide binding experiments.

## Author Contributions

GH, JK, SB and CZ measured and analyzed FRET samples. FSW performed peptide binding experiments. FD and NS created the PSG construct for the DMD simulations. NS performed and analyzed the DMD simulations. FSW, YH and FW performed and analyzed DS mapping experiments. FW designed expression constructs and prepared protein samples. CS and HS supervised confocal FRET experiments. MEB, HS, FD, NS, GH, and SB wrote the paper.

---

**Figure 1–figure supplement 1.**
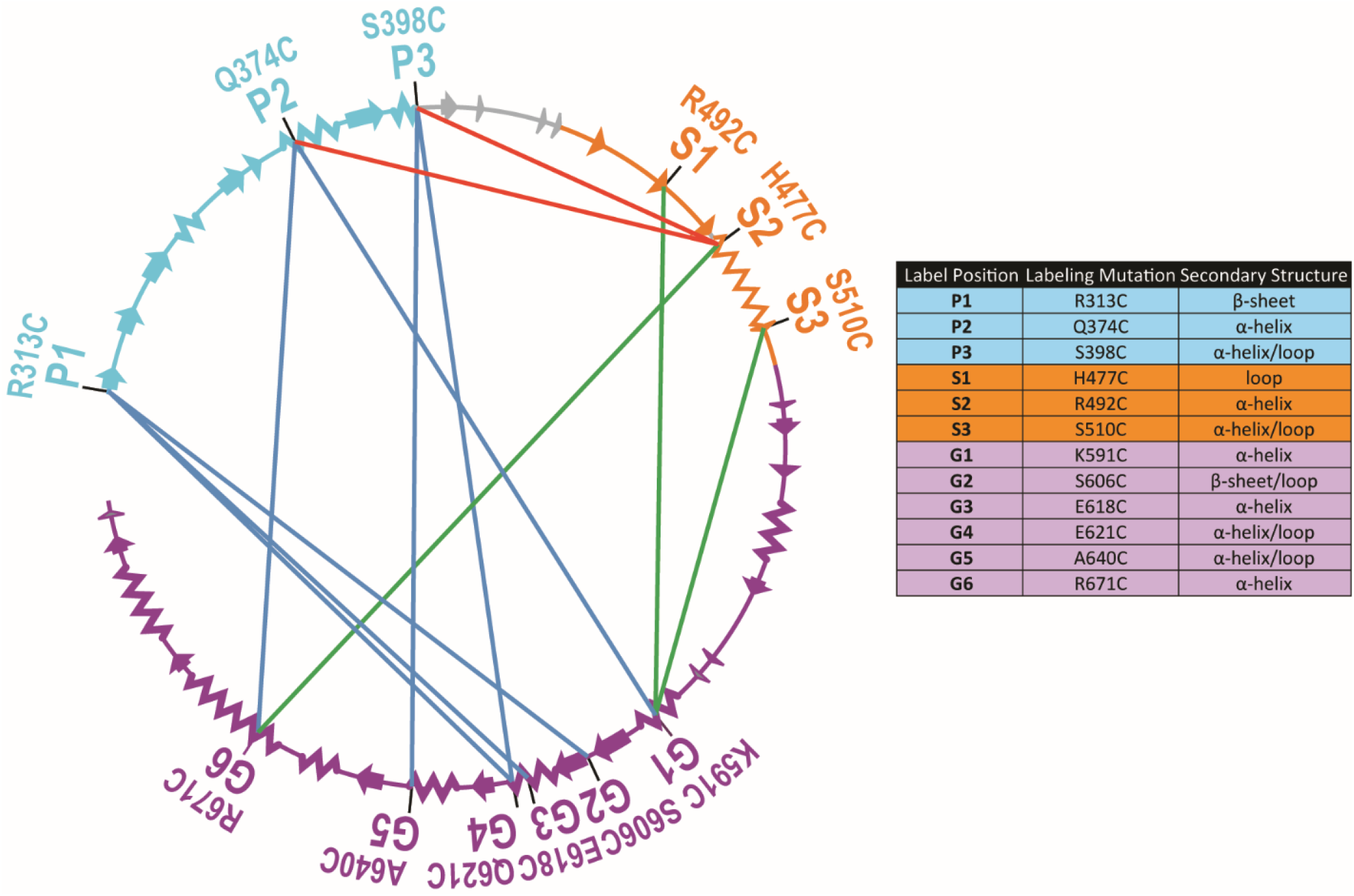
Connectivity in the FRET Network and Labeling Site Environments. The primary sequence of PSD-95 is shown in a circular representation with each domain colored: PDZ3 (cyan), SH3 (orange) and GuK (purple). The secondary structural elements are indicated: α helices (zig zag) and β sheets (arrows). The position of each labeling site is indicated by the domain and order in the primary sequence. Specific details for each labeling site are also provided in the associated table. The FRET pairs used for measurements are indicated by lines connecting the labeling sites used in that variant with FRET pairs spanning PDZ3-GuK (blue), PDZ3-SH3 (red), and SH3-GuK (green)

**Figure 1–figure supplement 2.**
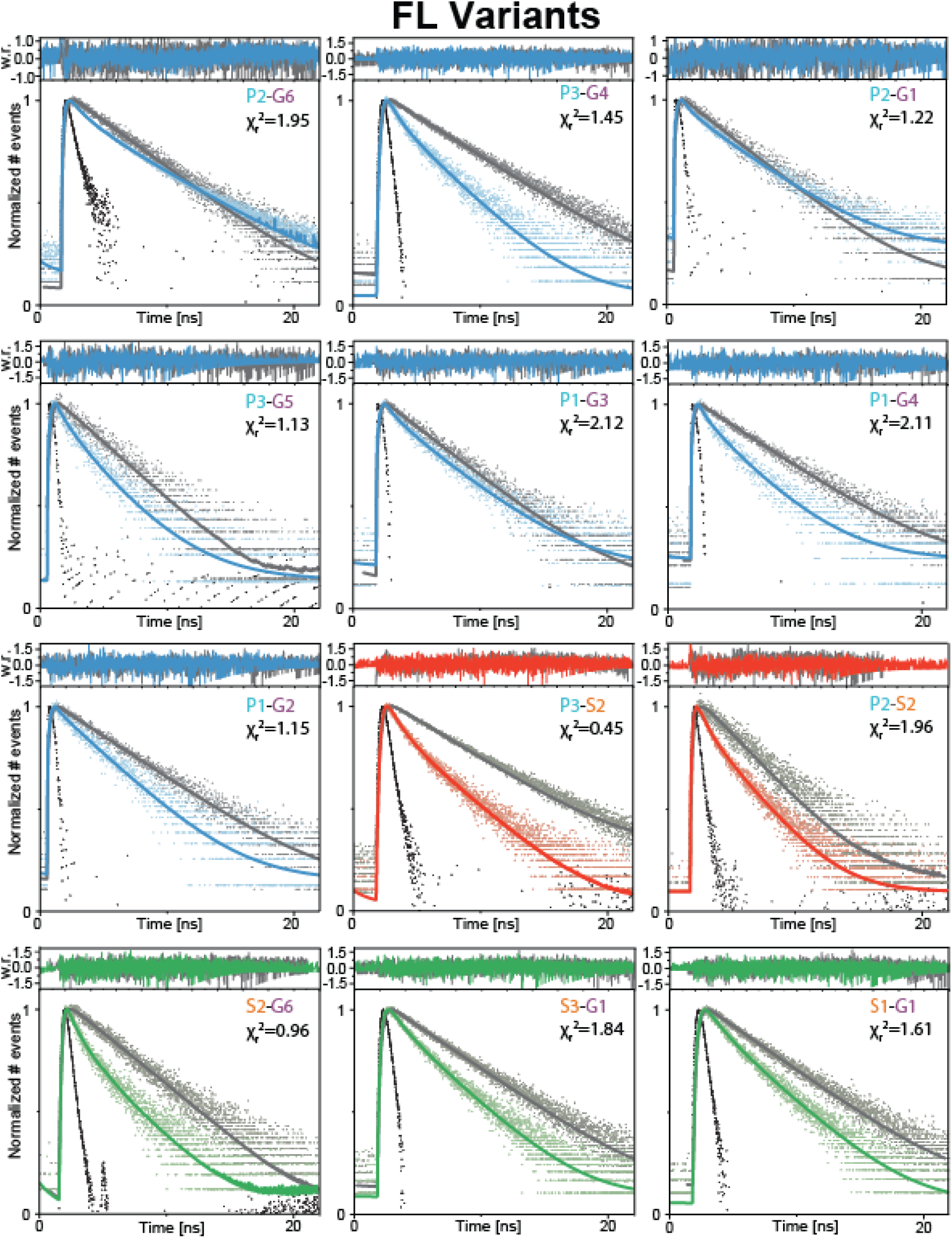
Global Fit of seTCSPC FRET-Sensitized Donor Fluorescence Decay Curves for Full Length Variants. Sub-ensemble Time Correlated Single Photon Counting decays for FRET pairs spanning PDZ3-GuK (blue), PDZ3-SH3 (red), and SH3-GuK (green). The instrument response function is shown in black, and the Donor-Only fluorescence decay is shown in gray. Raw histogram data are shown as points, with fits overlaid as lines. FRET results in quenching of the donor fluorescence and thus reduced fluorescence decay lifetime, and presence of more than one underlying population with different FRET results in multi-exponential fluorescence decays. Fit parameters can be found in Supplementary Files 1C and 1D. Details of the model and fit can be found in *Materials and Methods*.

**Figure 1–figure supplement 3.**
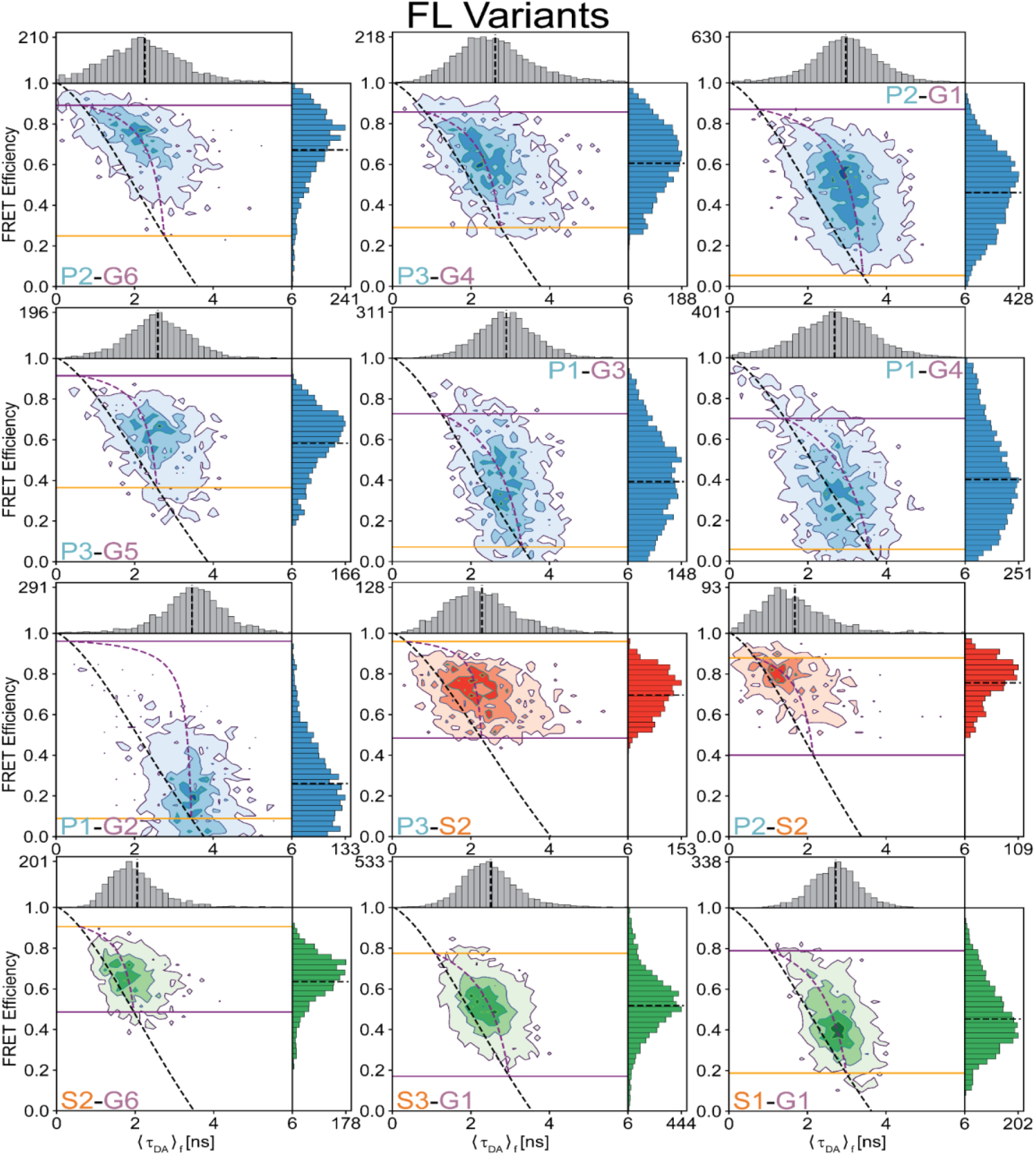
Multiparameter Fluorescence Histograms of Full-Length PSD-95 FRET Variants. MFD Histograms for FRET pairs spanning PDZ3-GuK (blue), PDZ3-SH3 (red), and SH3-GuK (green) as noted in each panel. Variant details can be found in Figure S1. Overlaid on the contour plots are the static FRET-lines (black, dashed), dynamic FRET-lines (purple, dashed), and solid horizontal lines corresponding to the limiting states A (orange) and B (purple) from seTCSPC. Also given are the burst-wise average values for the mean donor fluorescence lifetime (⟨⟨*τ*_*DA*_⟩_*f*_⟩) and mean FRET Efficiency (⟨*E*⟩) (black, dashed lines in 2D histograms). Dynamic exchange is immediately evident from broadening and skew rightward from the static FRET-line for each variant. Correction parameters for these histograms can be found in Supplementary Files 1A and 1B while parameters for the static and dynamic FRET-lines can be found in Supplementary Files 1F and 1G, respectively.

**Figure 2–figure supplement 1.**
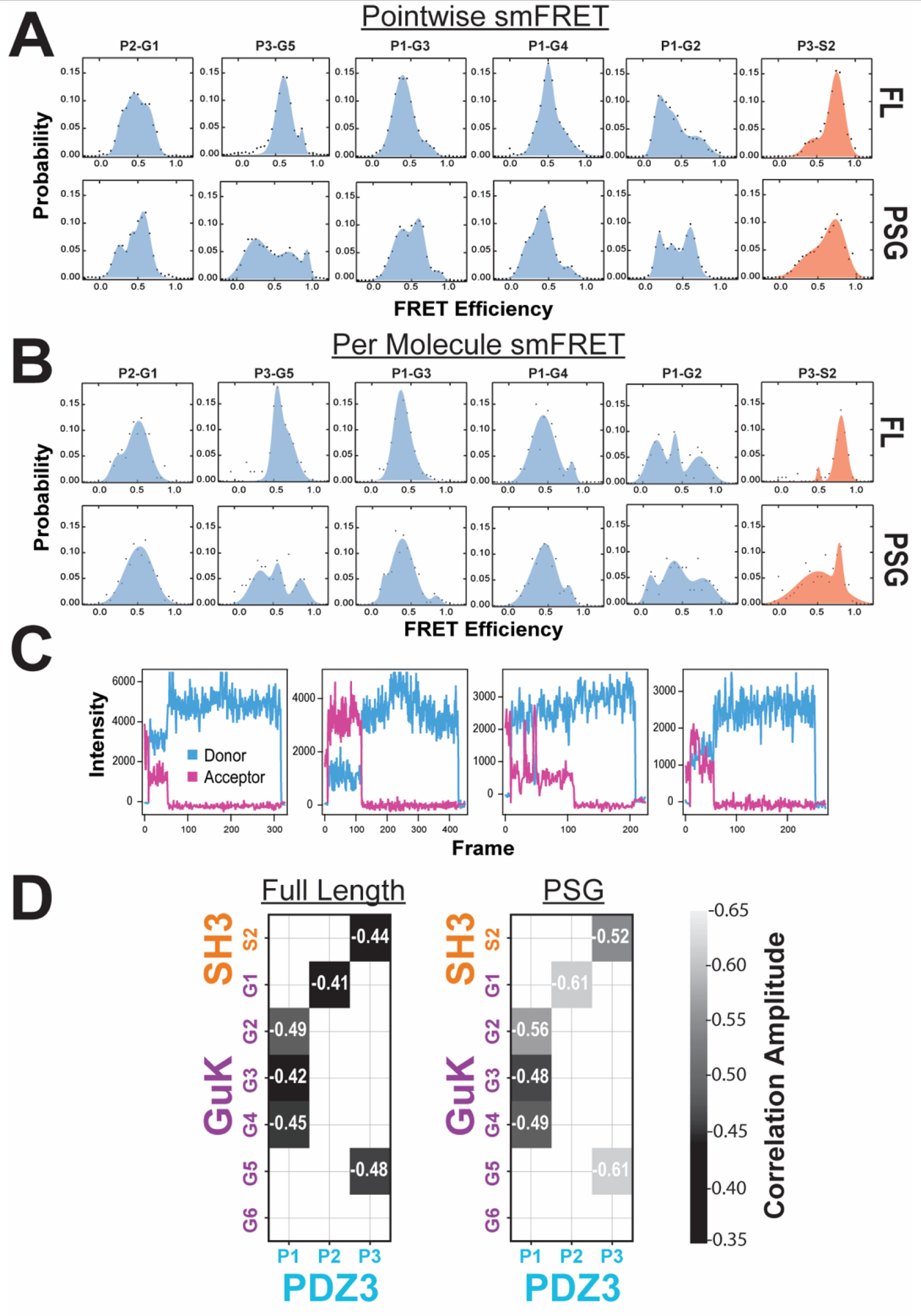
Effect of Truncation on PSD-95 as Measured with smTIRF. **A)** Population histograms of pointwise smTIRF data (per 100 millisecond frame) for variants with FRET pairs spanning PDZ3-GuK (blue), PDZ3-SH3 (red) as indicated above each panel. Shown are data for full-length PSD-95 (top) and the truncated PSG (bottom). **B)** Time-averaged (per molecule) population histograms from the same smTIRF data shown in panel A with identical coloring. **C)** Representative single-molecule time traces for variant P2-G1 in the PSG truncation showing static and dynamic molecules. **D)** Matrix representation of site-specific dynamics in full-length PSD-95 (left) and the PSG truncation (right). The axes specify the domains and labeling sites with each variant placed at the intersection between sites used. Shown are donor-acceptor cross-correlation amplitudes as calculated from smTIRF time traces before photobleaching occurred. The cross-correlation amplitude captures anticorrelated FRET transitions on the sub-second timescale such as those visible in panel C.

**Figure 2–figure supplement 2.**
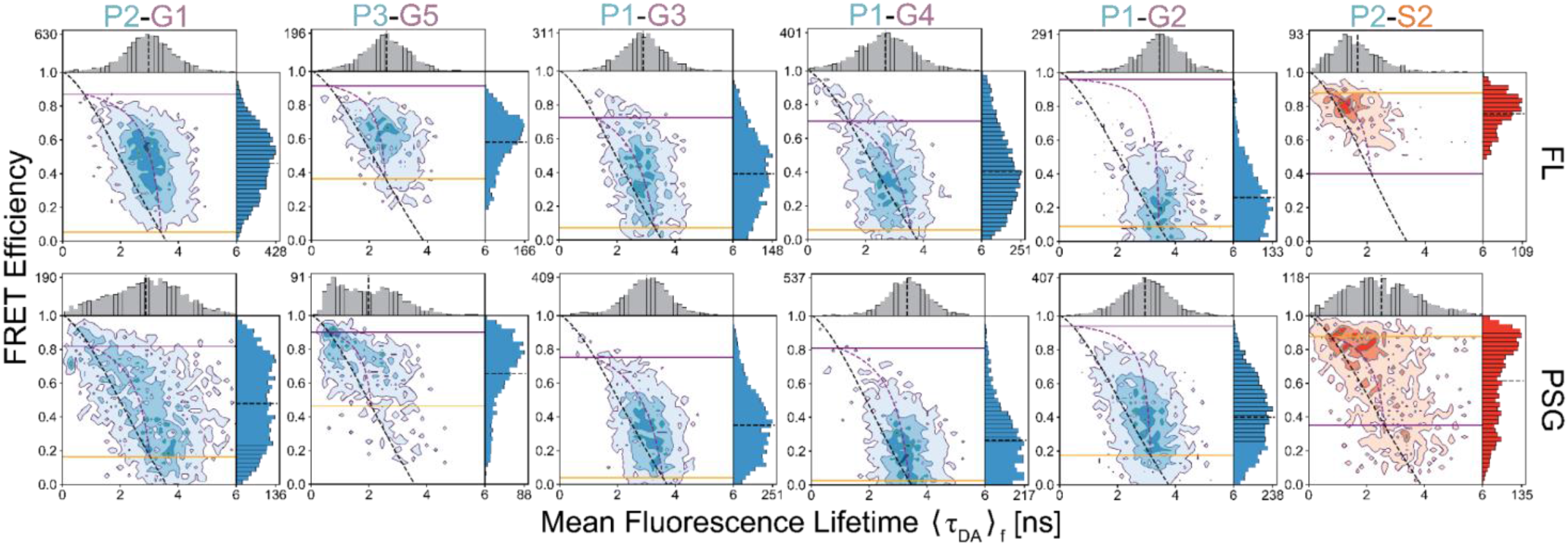
Comparison of Multiparameter Fluorescence Histograms of PSG-Truncated and Full-Length PSD-95 FRET Variants. MFD plots of PSG truncations (bottom). FL variants with same labeling sites are shown for comparison (top). The labeling sites are indicated within each panel. Dynamic lines and limiting states for PSG variants were determined from global fitting of donor fluorescence decays of PSG variants shown in Figure S5. Experimental parameters can be found in Supplementary Files 1A, and 1B. Fit parameters for determination of the limiting states can be found in Supplementary Files 1C-E. Parameters for the calculation of static and dynamic FRET lines can be found in Supplementary Files 1F and 1G.

**Figure 2–figure supplement 3.**
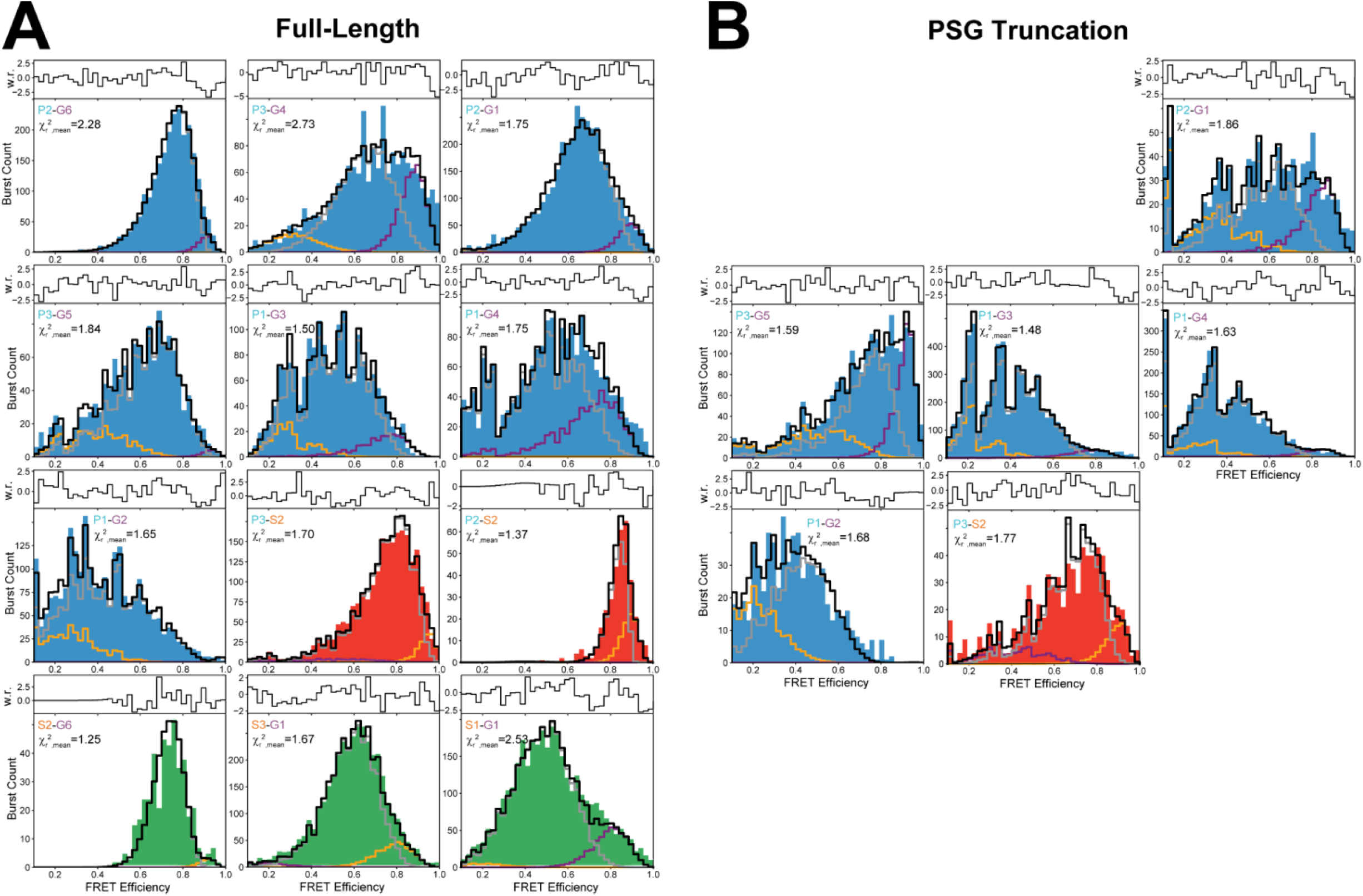
Photon Distribution Analysis Histograms. Representative photon distribution analysis for both Full Length PSD-95 and PSG truncated variants. Data shown is binned using 2 ms time windows. Fits were performed globally with 1 ms, 2 ms, and 3 ms time windows for each sample. The displayed 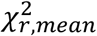 is the simple mean of 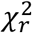 for fits of all three time windows. Raw data is color coded as blue (PDZ3-GuK), red (PDZ3-SH3), or green (SH3-GuK). Data is binned using 50 evenly spaced bins in FRET efficiency. Fits are shown in grey (dynamic fraction), orange (state A static fraction), purple (state B static fraction), and black (sum of three components). Fit parameters can be found in Supplementary File 2.

**Figure 2–figure supplement 4.**
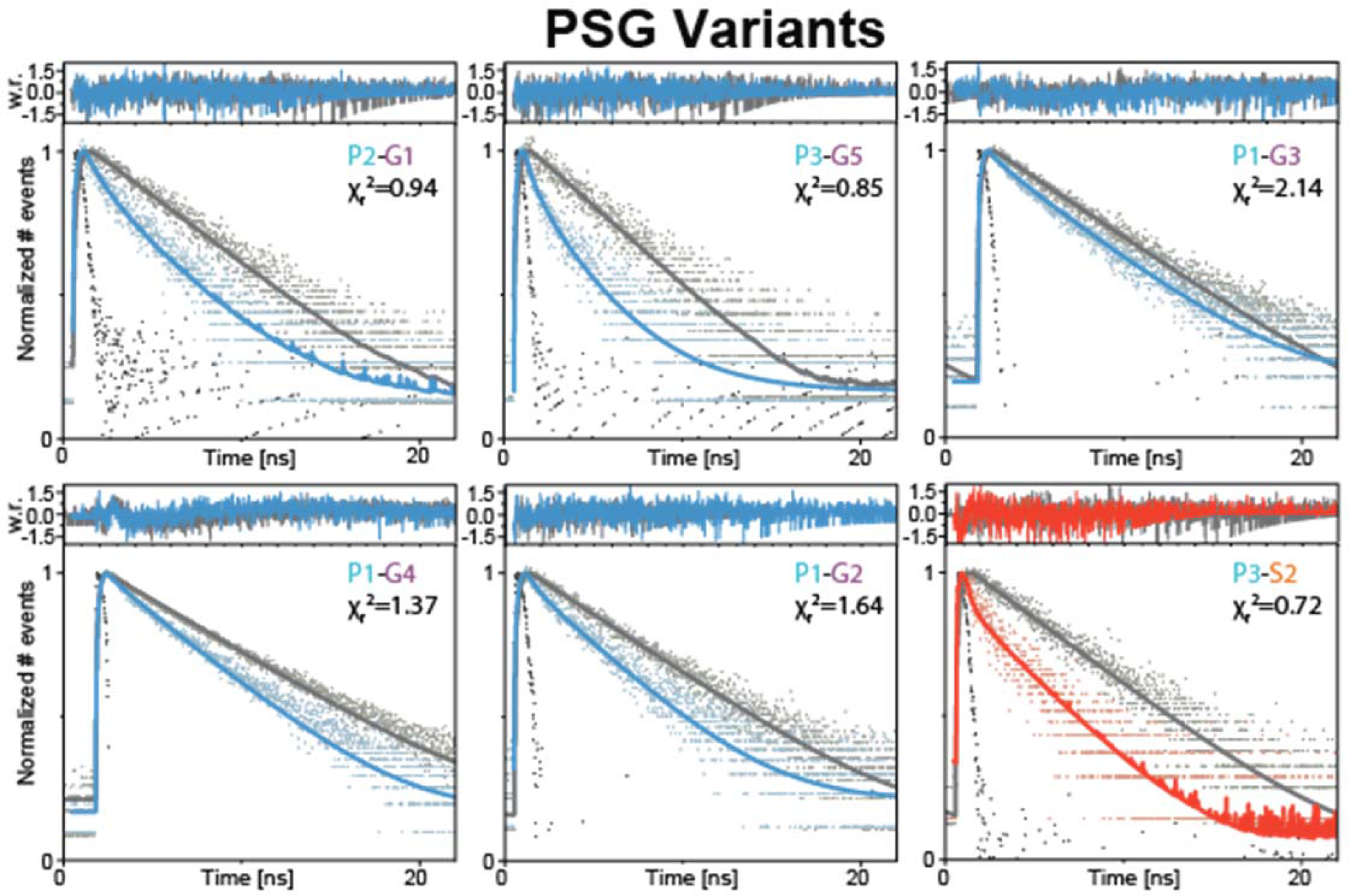
Global Fit of seTCSPC FRET-Sensitized Donor Fluorescence Decay Curves for Truncated PSG Variants. Fit parameters can be found in Supplementary Files 1C and 1D. Fits are analogous to those shown in Figure S2 but for PSG variants Donor only samples colored grey. FRET-sensitized donor decays colored blue for PDZ3-GuK; red for PDZ3-SH3.

**Figure 3–figure supplement 1.**
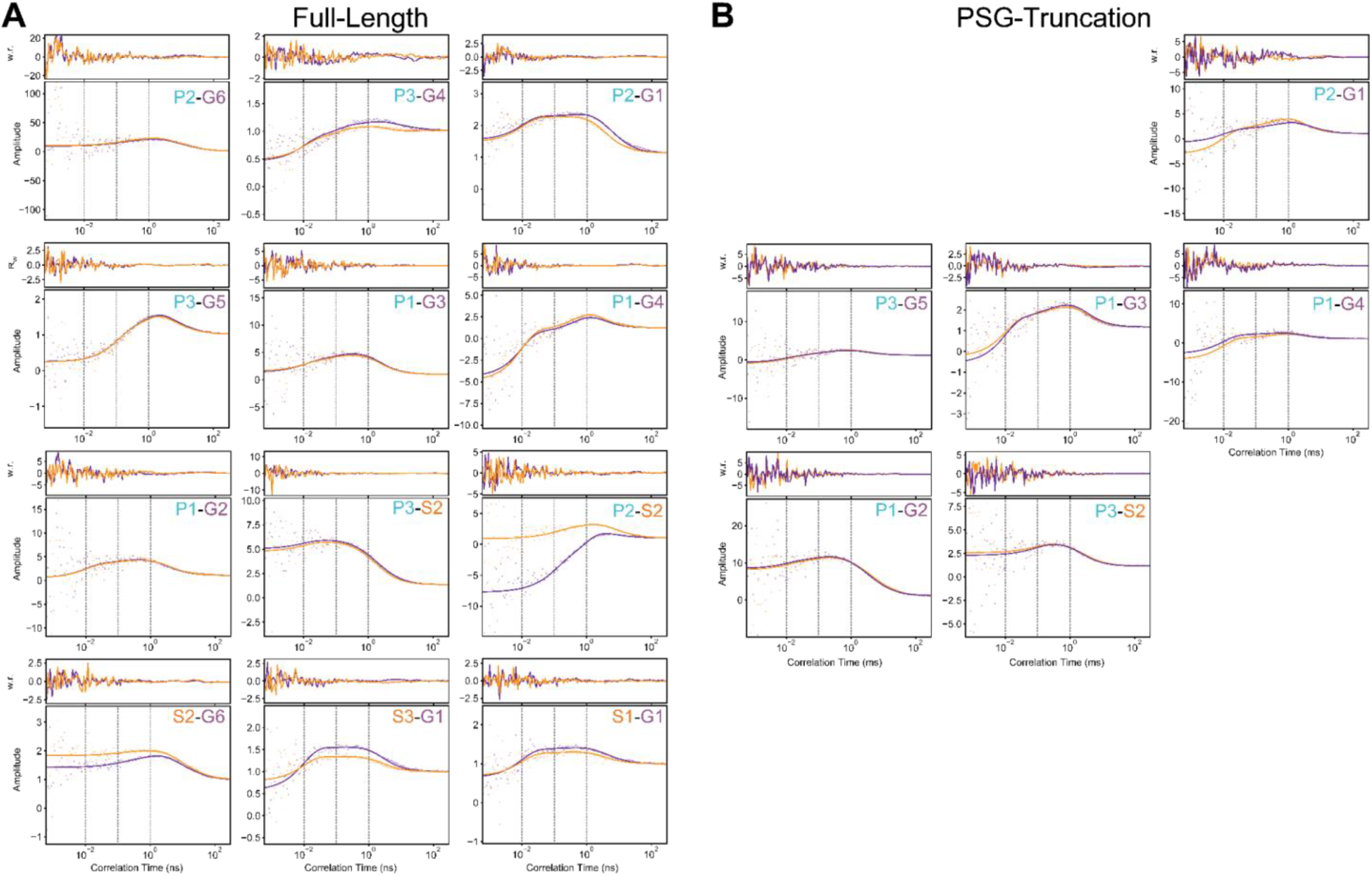
Filtered Fluorescence Correlation Spectroscopy Fits. **A)** Cross-correlation curves for full-length PSD-95 variants as indicated within each panel. **B)** Cross-correlation curves for truncated PSG variants as indicated within each panel. Curves were fit with diffusion timescales fit based on autocorrelation curves. Cross-correlation decay timescales are fixed to .01 ms, .10 ms, and 1.00 ms (gray, vertical dashes) such that relative amplitudes corresponding to each timescale can be directly compared (Figure 3). Weighted residuals are displayed in upper panels. Raw correlation data is shown as points with lines overlaid for fit curves. Purple data corresponds to low-FRET to high-FRET component cross-correlation, while orange corresponds to low-FRET to high-FRET. Individual amplitudes are shared between the datasets, but each component is allowed a global scaling factor (Eqns. S2 and S3). Details of the model functions are available in *Materials and Methods*.

**Figure 3–figure supplement 2.**
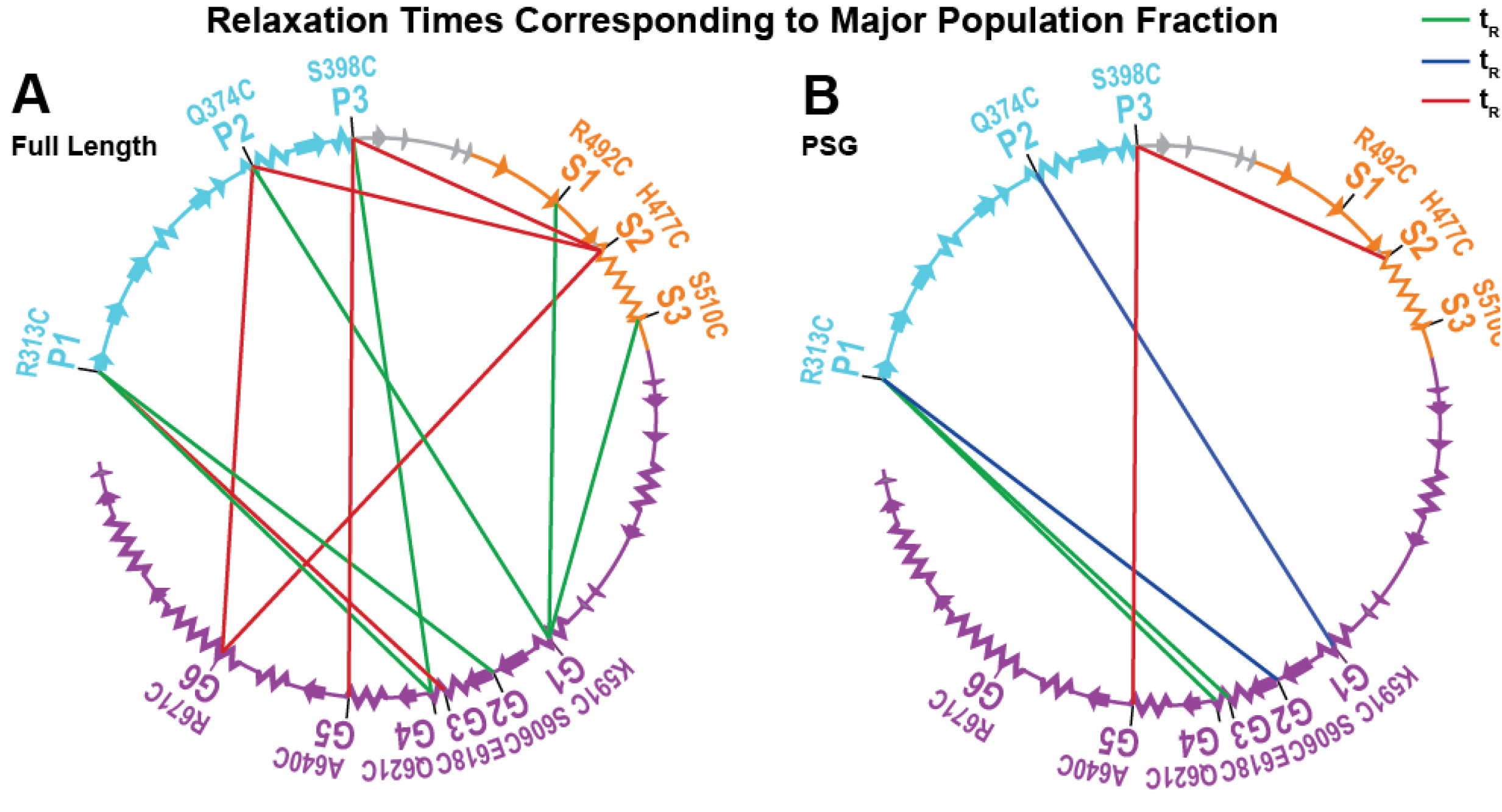
Summary of Dynamics Probed by fFCS Mapped to the Primary Sequence. Cartoon representations of the sequence of the PSG of PSD-95, with coloring and secondary structure as in Figure 1. Lines connecting the fluorophore labeling sites are colored according to whichever relaxation time from global fFCS analysis corresponds to the major population fraction for each site (magenta, t_R1_ = 0.01 ms, green, t_R2_ = 0.10 ms, and dark blue, t_R3_ = 1.00 ms). The most dominant relaxation times, t_R1_ and t_R3_ correspond to fast local motions and domain-scale conformational exchange, respectively. Summary diagrams are provide for both **A)** full length and **B)** PSG truncated variants.

**Figure 4–figure supplement 1.**
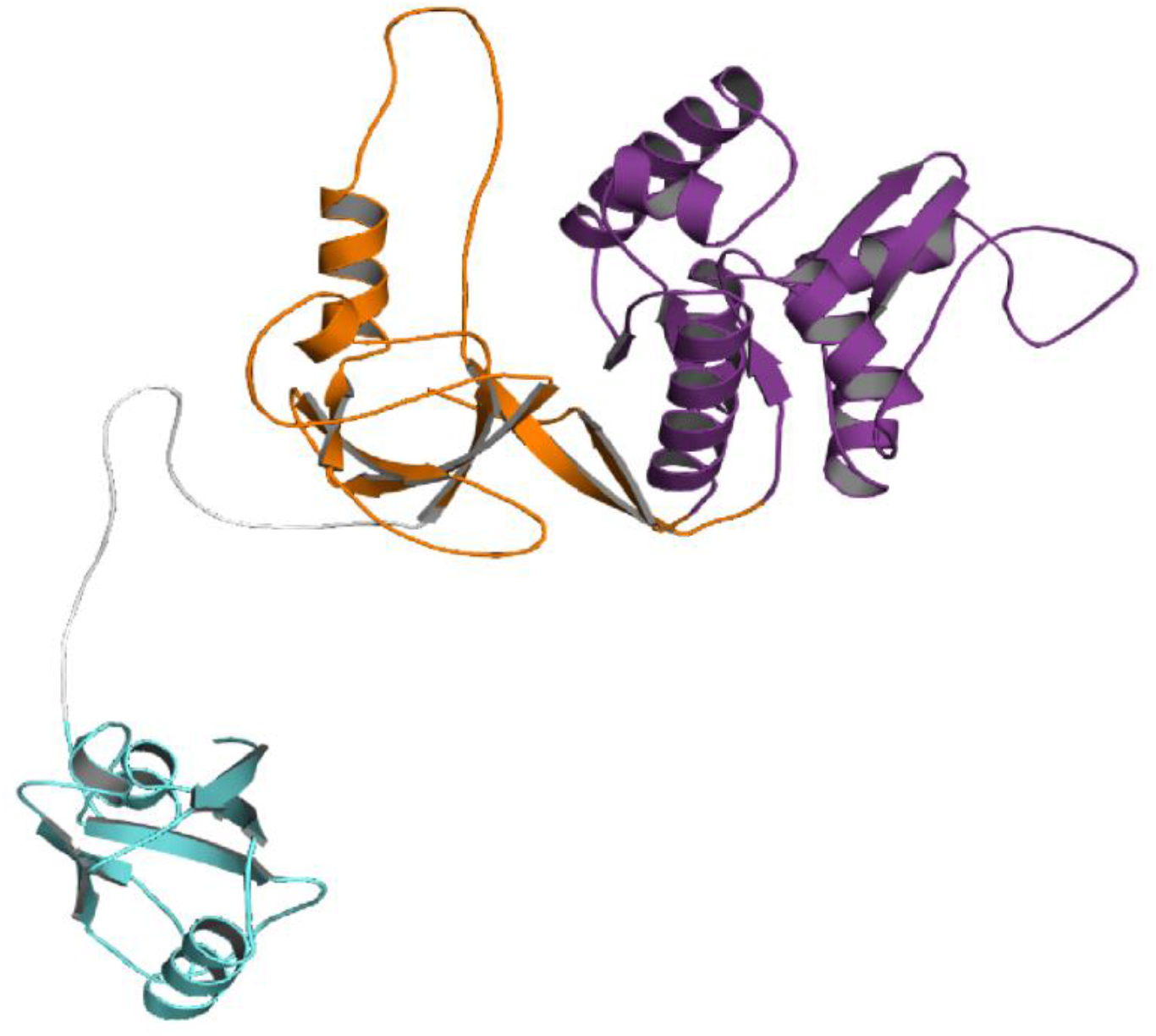
The Starting Conformation for the PSG Supramodule from PSD-95 Used in DMD Simulations. Cartoon representation of the initial model for DMD simulations showing PDZ3 (cyan), SH3 (orange) and GuK (purple). To avoid biasing the interactions, PDZ3 was positioned away from the SH3-GuK without any contacts. The model was constructed from the crystal structures of PDZ3 (1TP5) and SH3-GuK (1KJW). PDZ3 was placed in a random orientation without interdomain contacts to avoid bias. The PDZ3-SH3 linker and missing loops were reconstructed using our in-house loop reconstruction program ‘medusa-loop’.

**Figure 4–figure supplement 2.**
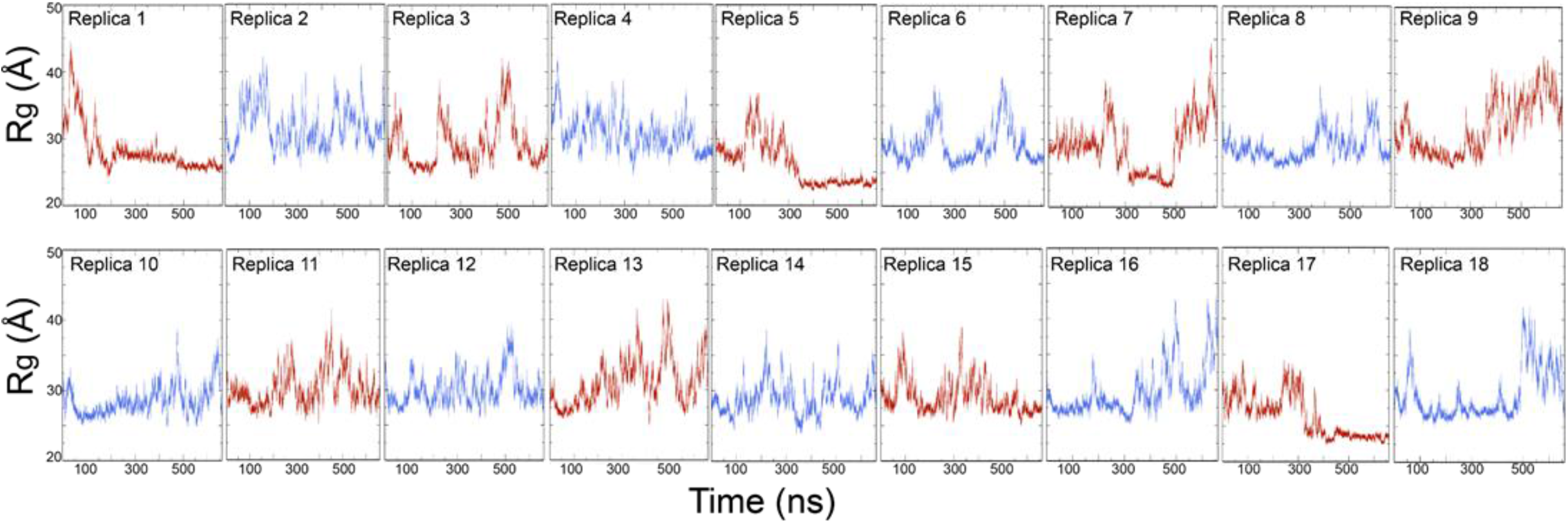
Time evolution of the radius of gyration (*R*_*g*_) of PSG supramodule for 18 replicas DMD simulations. The total simulation time for each replica is 660 ns and amounts to a total simulation time of 11.9 μs. We used the last 400 ns of replica exchange trajectories for the statistical analysis.

**Figure 4–figure supplement 3.**
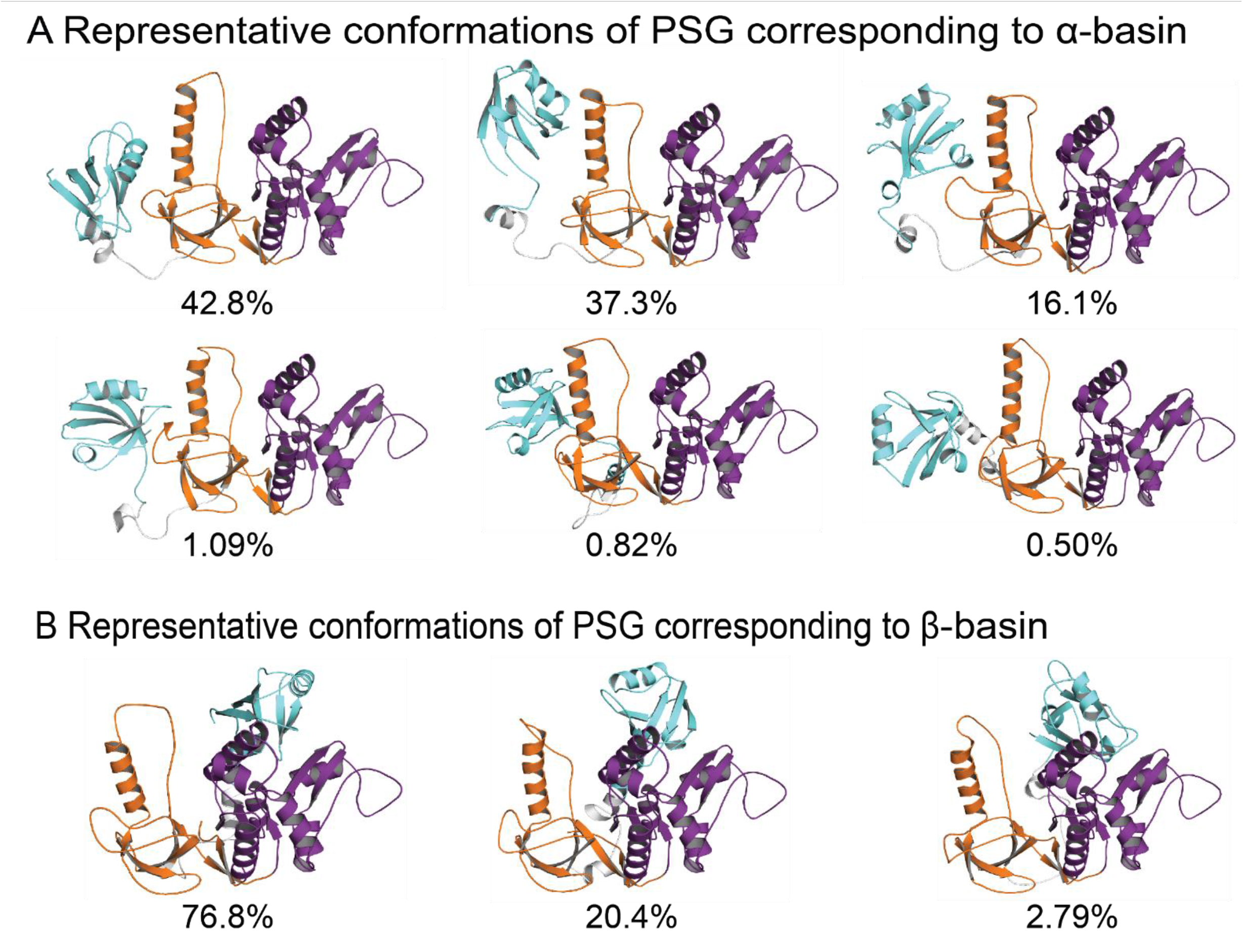
Representative Conformations of PSG Supramodule from PSD-95. **A)** Representative conformations and their population occupancy in the highly sampled α-basin with PDZ3 interacting with the SH3 domain. A multiplicity of states is sampled within the fuzzy α-basin. The top centroid cluster in the α-basin shows PDZ3 to reorient away from the HOOK helix and sample the β1-β2 loop (RT loop in canonical SH3 domain). The pairwise contact maps from DMD show that interactions within the α-basin involved degenerate electrostatic interactions engaged by all conformations with occasional hydrophobic interactions leading an occluded PDZ3 binding pocket. **B)** Representative conformations and their population in the β-basin with PDZ3 interacting with the GuK domain of PSG. The β-basin was comparatively well defined due to involvement of exposed hydrophobic residues in β3-α1 of PDZ3 and a relatively hydrophobic surface in GuK formed by α7, α5, and β10-11 (e.g., L349-F684, L342-L608), which support a limited range of conformations. Both basins infrequently engaged the canonical GLGF motif, which would prevent peptide binding to PDZ3. An RMSD cutoff of 8.0 Å was used to select the centroid clusters of α- and β-basins. PDZ3 is colored cyan, SH3 colored orange, and GuK colored purple.

**Figure 4–figure supplement 4.**
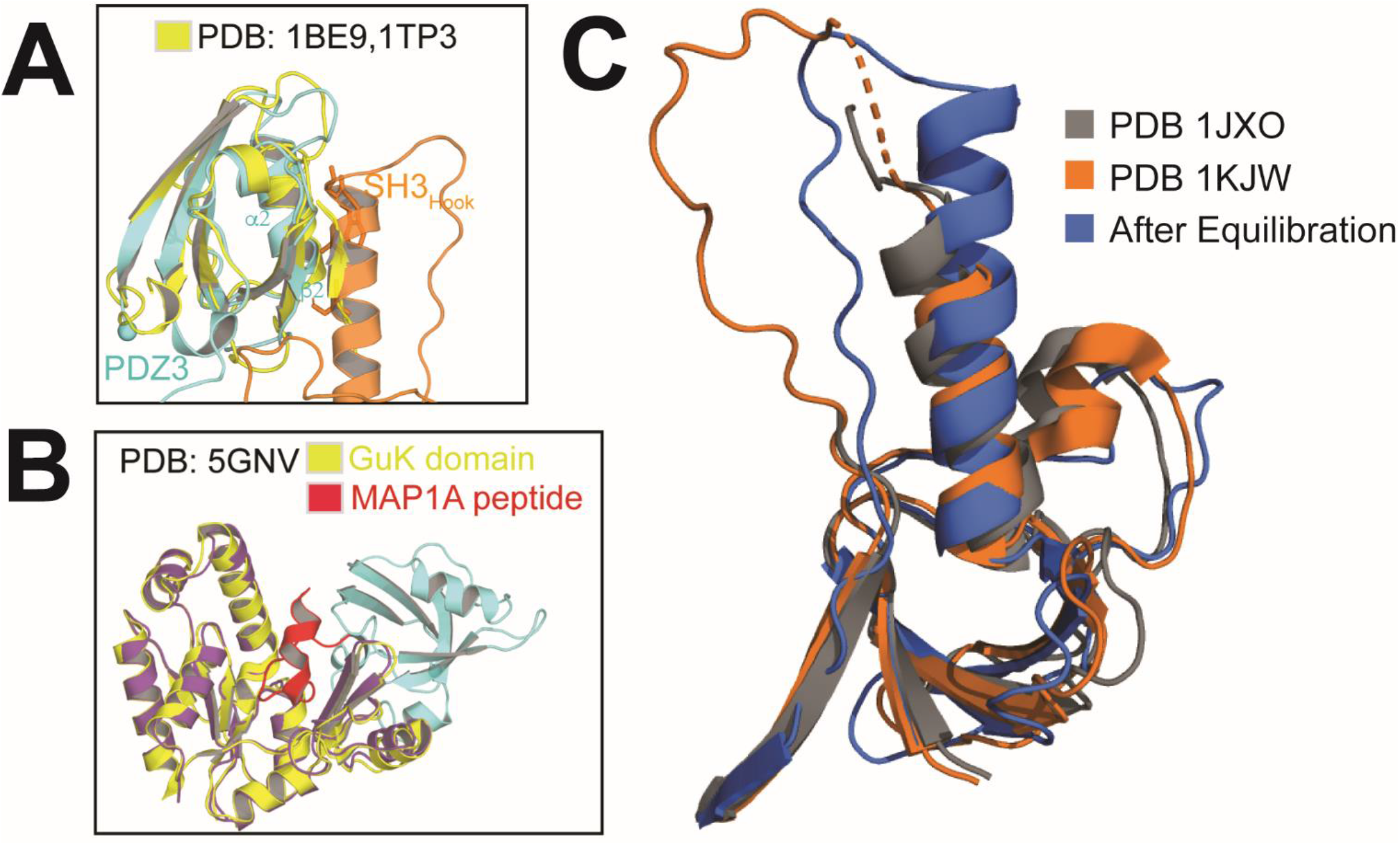
Comparison of Equilibrated Conformations Observed in DMD to Published Crystal Structures. **A)** Steric occlusion of the PDZ3 ligand-binding pocket within the α-basin. The structure of PDZ3 bound to a short peptide (1TP3, yellow) is aligned with PDZ3 from a representative α-basin model (cyan). The ligand is shown as a beta strand (yellow) that overlaps with the SH3 HOOK insertion (orange). **B)** Lack of steric occlusion in the β-basin. The structure of GuK (yellow) bound to a MAP1A peptide (red) is aligned with a representative β-basin model (purple). The canonical ligand binding pockets of GuK and PDZ3 remain accessible in the β-basin.

**Figure 5–figure supplement 1.**
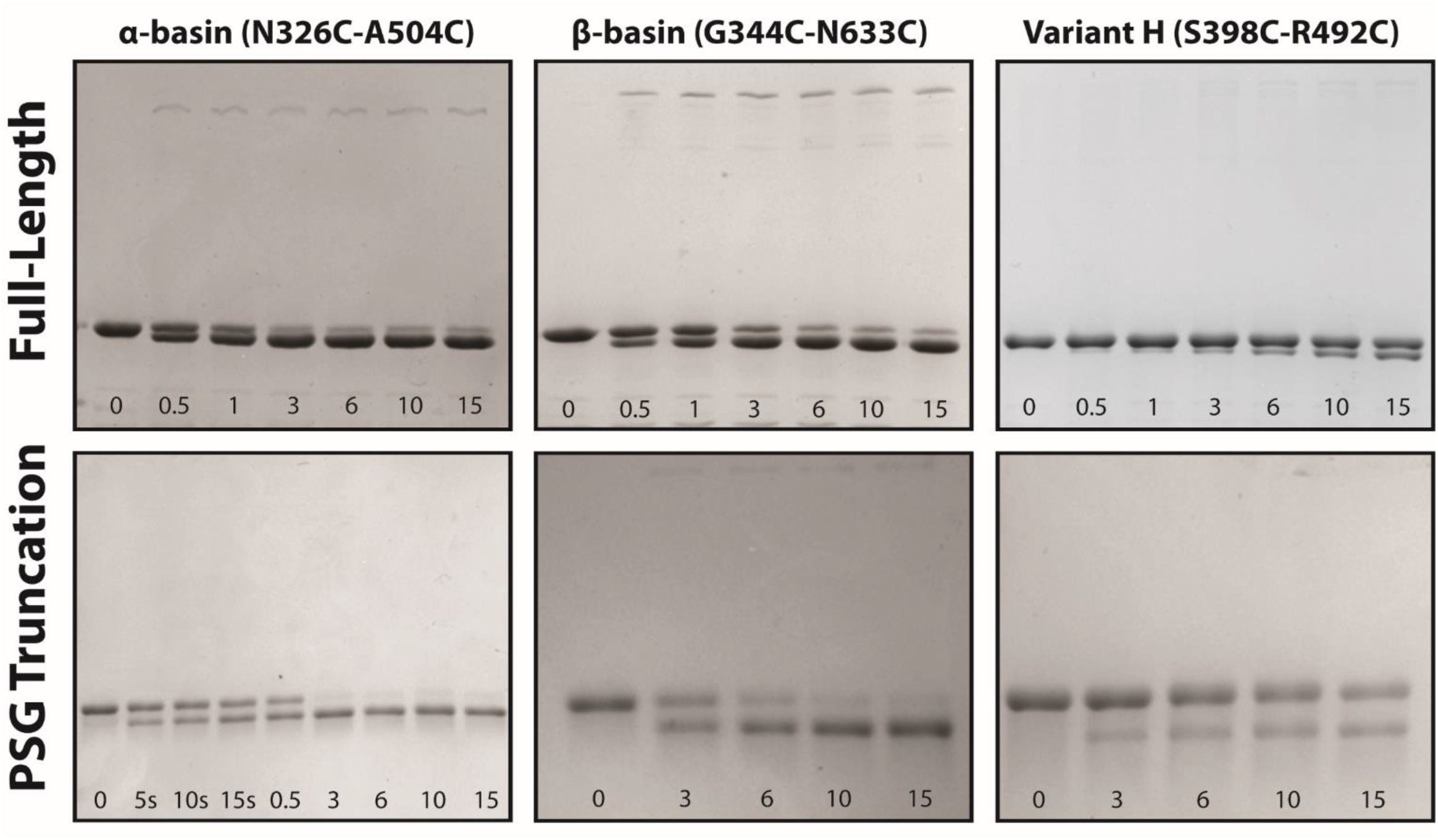
Disulfide Mapping of the Contact Interfaces from DMD. Representative non-reducing SDS-PAGE gels used to extract kinetic information about the efficiency and rates of disulfide formation. The top row shows 7.5% acrylamide gels for full-length PSD-95 variants. The bottom row shows 10% acrylamide gels used for PSG truncations. The variant and the introduced mutations are noted above each column. The time for each lane is indicated within each panel in minutes unless indicated otherwise. The background-subtracted intensity of each band was obtained using ImageJ to calculate the background-subtracted mean intensity, which are plotted in Figure 5. Each experiment was repeated in triplicate.

**Figure 6–figure supplement 1.**
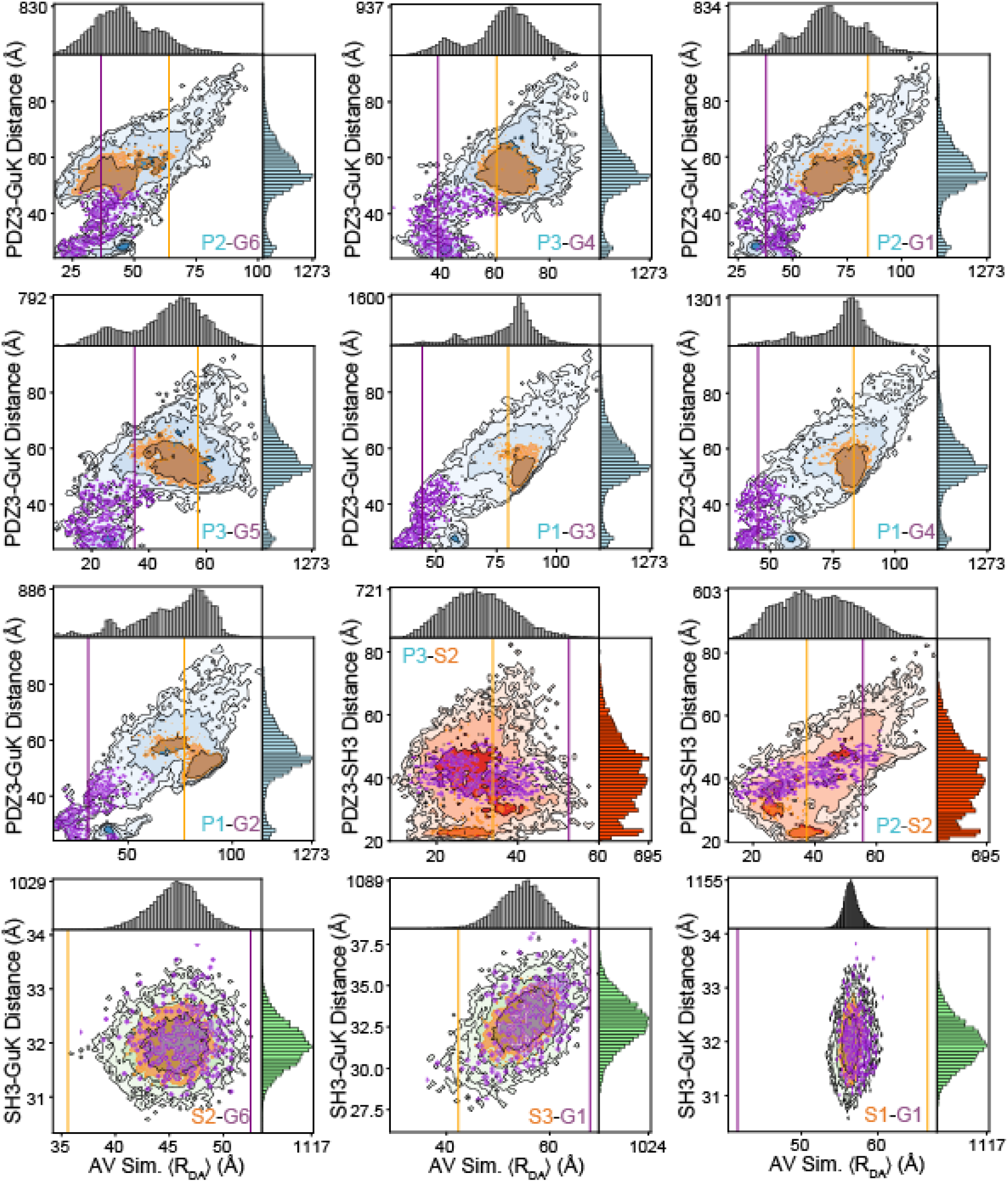
Responsiveness of Individual Variants to the Underlying Conformational Distribution. Shown are the simulated interdye distances ⟨*R*_*DA*_⟩ from accessible volume (AV) simulations using DMD structures. The 20871 snapshot structures from DMD were used to calculate the accessible volumes for the dyes at each labeling position, which were then used to calculate average interdye distances for each snapshot structure. These interdye distances for each variant are plotted against the distance between centers of mass (CoM) for the relevant domains. These 2D plots provide a qualitative analysis of how each FRET pair reflects changes in the underlying conformation. The limiting state distances for each variant are shown as vertical lines for state A (orange) and state B (purple). Overlaid are contours corresponding to DMD structures residing in state A (orange) and state B (purple), which are within the distance uncertainty from sub-sampling analysis of TCSPC fluorescence decays and the AV simulation distances. For most variants, limiting state distances qualitatively agree with the locations of local maxima in the simulated interdye distance distributions. However, some variants show both maxima along the same vertical line indicating that the conformational dynamics are not fully captured by that FRET pair. Further, presence of FRET pairs spanning SH3-GuK show the most difference between simulated and measured distances, likely owing to the limited dynamics of these domains *in silico*.

**Figure 6–figure supplement 2.**
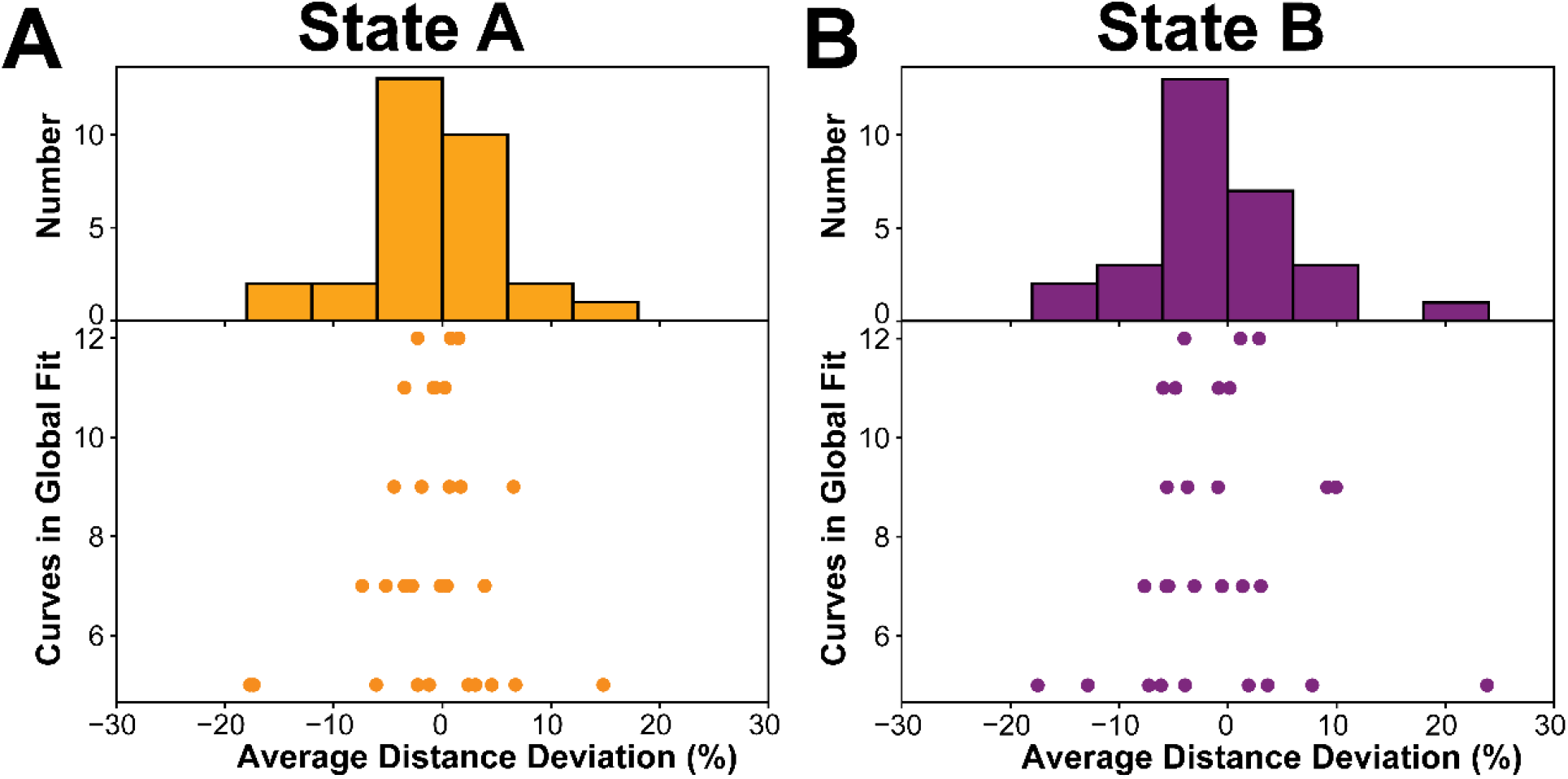
Robustness of the FRET Network to Sub-sampled Global Fitting. To quantify the robustness of our global fit to the seTCSPC fluorescence decays for our FRET network, we randomly selected sub-samples of the FRET network in groups of five, seven, nine, eleven, and twelve variants and repeated the global fitting on the reduced FRET network. Subsets were generated such that each variant was included in at least three subsets to avoid underrepresentation of individual variants. Each subset was re-analyzed in Chisurf using a two-state model with distances free to vary. Percent errors for each fit were calculated relative to distances from the global fit of the full FRET network, which is summarized for all fits in the associated histogram. Distance errors are reported separately for each state from the two-state model. Distances and associated widths resulting from subsampled global fitting are summarized in Supplementary File 6B. **A)** Average distance deviation for state A from global fitting of individual sub-sampled FRET networks. **B)** Average distance deviation for state B. The histograms show that the distributions are centered on the reported distances regardless of the number of variants. These summary statistics indicate convergence toward the center of the distribution as the number of samples globally fit is increased.

**Figure 6–figure supplement 3.**
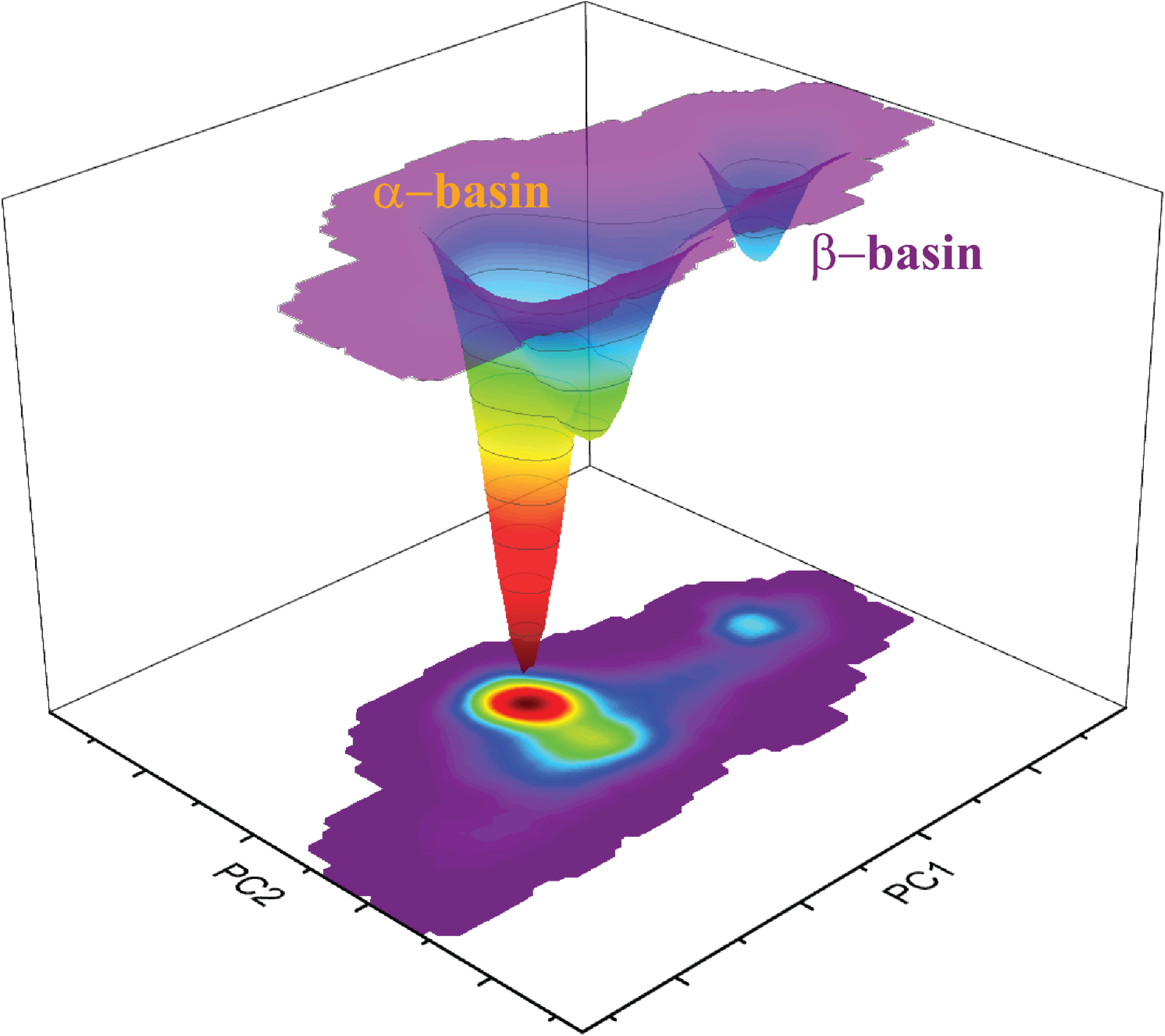
Principal Component Analysis of COM and AV Data. In order to visualize the conformational landscape of the PSG, we performed principal component analysis using the COMs distances between domains and the simulated ⟨*R*_*DA*_⟩ of PDZ3 to SH3 and GuK as shown in Figure 6-Supplementary File 1. Data were analyzed using scikit-learn (sklearn) in Python. Data were first standardized such that each set of inter-COM and AV-derived interdye distances had mean 0 and unit variance using the sklearn.preprocessing.StandardScaler. Principal components were calculated using sklearn.decomposition.PCA with two components. The surface depth corresponds to the relative number of structures from DMD simulations which occupy some region of the principal component (PC) space. PCA results in basins α and β separated along PC1, which explains 65% of the standardized dataset variance. The variance along PC2 (16% explained variance ratio) mostly corresponds to apparent heterogeneity in basin α. Additionally, there is an apparent saddle point corresponding to the transition pathway between the two basins along PC1. This suggests that the motions corresponding to heterogeneity in basin α are distinct from those corresponding to interdomain transitions as observed by FRET. This surface was rescaled for Figure 6 such that the integrated volume of the two basins were equivalent to the population fractions for states A and B obtained from TCSPC analysis.

**Figure 7–figure supplement 1.**
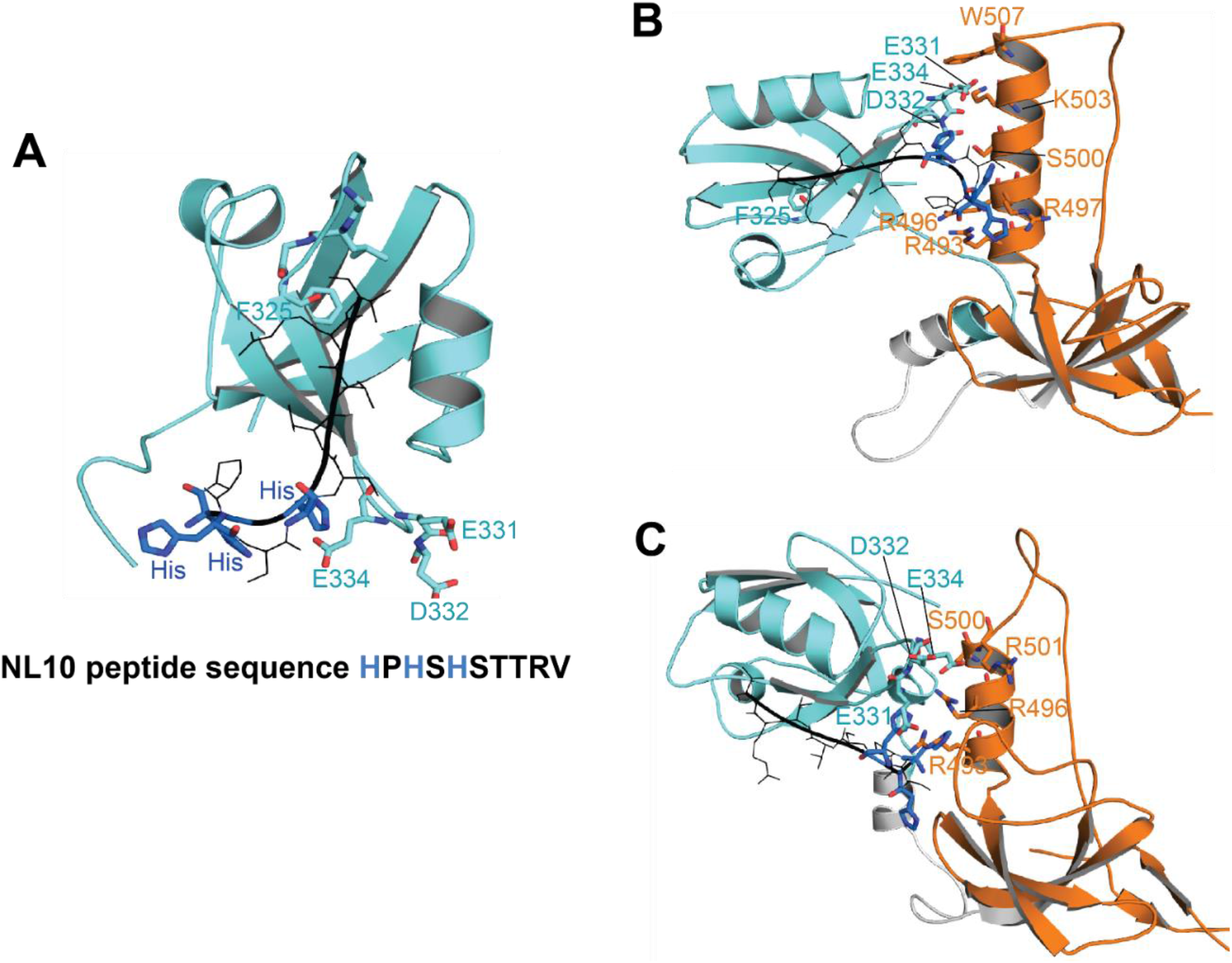
Docking of Neuroligin to PDZ3 and the PSG Supramodule in α-basin. **(A)** Docked conformation of NL10 peptide bound to the PDZ3. The NL10 peptide is shown in black with the three histidine residues shown in blue. When protonated, these histidine residues extend towards E331, D332, and E334 in PDZ3, which may explain the higher binding affinity at low pH. At physiological pH, the negatively charged residues will inhibit binding of uncharged histidines. **(B-C)** Representative conformations of NL10 peptide binding to PDZ3 within the PSG α-basin. In the context of PSG, positively charged residues in the SH3 HOOK insertion form salt-bridges with the negatively charged residues in PDZ3. These electrostatic interactions sequester these negative charges to stabilize the binding of NL10 at physiological pH.

